# Multiple overlapping hypothalamus-brainstem circuits drive rapid threat avoidance

**DOI:** 10.1101/745075

**Authors:** Matthew Lovett-Barron, Ritchie Chen, Susanna Bradbury, Aaron S Andalman, Mahendra Wagle, Su Guo, Karl Deisseroth

## Abstract

Animals survive environmental challenges by adapting their physiology and behavior through homeostatic regulatory processes, mediated in part by specific neuropeptide release from the hypothalamus. Animals can also avoid environmental stressors within seconds, a fast behavioral adaptation for which hypothalamic involvement is not established. Using brain-wide neural activity imaging in behaving zebrafish, here we find that hypothalamic neurons are rapidly engaged during common avoidance responses elicited by various environmental stressors. By developing methods to register cellular-resolution neural dynamics to multiplexed *in situ* gene expression, we find that each category of stressor recruits similar combinations of multiple peptidergic cell types in the hypothalamus. Anatomical analysis and functional manipulations demonstrate that these diverse cell types play shared roles in behavior, are glutamatergic, and converge upon spinal-projecting brainstem neurons required for avoidance. These data demonstrate that hypothalamic neural populations, classically associated with slow and specific homeostatic adaptations, also together give rise to fast and generalized avoidance behavior.

---

Environmental changes prompt organisms to adapt their physiology and behavior to new circumstances, responses that promote survival in a changing world. These adaptive responses to environmental stressors occur over multiple timescales, including rapid avoidance of threats within seconds of stressor detection, and slower homeostatic adaptations to these stressors on the timescale of minutes to hours or longer (Wingfield, 2003; Heinrichs & Koob, 2004; Dallman, 2005; Joëls & Baram, 2009). In vertebrates, slow homeostatic adaptations are largely mediated by the hypothalamus, an evolutionarily conserved set of structures that integrate internal need and sensory cues to coordinate appropriate behavioral, physiological, and endocrine responses (Ulrich-Lai & Herman, 2009; Sternson, 2013; Saper & Lowell, 2014; Herman & Tasker, 2016; Biran et al., 2018).

The hypothalamus is composed of a rich set of distinct cell types, often distinguished by the expression of specialized neuropeptide transmitters (Joëls & Baram, 2009; Herget & Ryu, 2015; Romanov et al., 2017; Moffitt et al., 2018; Romanov et al., 2019). Different homeostatic needs are known to engage specific peptide-releasing hypothalamic neurons to drive adaptive responses. For instance, blood or cerebrospinal fluid hyperosmolarity recruits neurons in the paraventricular hypothalamus that release arginine vasopressin or oxytocin, among other transmitters (McCormick & Bradshaw, 2006; Bourque, 2008; Zimmerman et al., 2017; McKinley et al., 2019). Energy imbalance recruits neurons in the paraventricular hypothalamus that release oxytocin, corticotrophin-releasing hormone, or thyrotropin-releasing hormone, as well as neurons in the arcuate hypothalamus that release neuropeptide-Y(Gao & Horvath, 2007; Aponte et al., 2011; Krashes et al., 2011; Hill, 2012; Sternson, 2013). Changes in blood or cerebrospinal fluid acidity recruit neurons in the lateral hypothalamus that release hypocretin/orexin or melanin-concentrating hormone, among other neurons (Williams et al., 2007; Burdakov et al., 2013). Changes in body temperature recruit neurons in the dorsomedial hypothalamus that release prolactin-releasing peptide, and neurons in the preoptic hypothalamus that release brain-derived neurotrophic factor and pituitary adenylate cyclase-activating polypeptide, among other cells (Morrison & Nakamura, 2011; Song et al., 2016; Tan et al., 2016; Tan & Knight, 2018). These distinct sets of peptidergic cell types initiate the specific behavioral and physiological adaptations associated with these homeostatic needs; such adaptations typically occur over the course of minutes to hours, initiated by the central release of neuropeptides and classical neurotransmitters, and/or systemic peptide release into the circulatory system.

In addition to slower adaptive responses, animals can also exhibit faster avoidance responses, often on the timescale of seconds (Wingfield, 2003). For instance, animals will avoid stimuli or environments with extreme temperatures (Tan et al., 2016), carbon dioxide (Spiacci Jr et al., 2018), or salinity (Oka et al., 2013). Avoidance behaviors are also observed in virtual states of hunger and thirst, evoked by experimental stimulation of hypothalamic neurons encoding these need states (Betley et al., 2015; Allen et al., 2017; Leib et al., 2017). If the option to escape is available, evasive behaviors can allow an animal to avoid the need to produce systemic homeostatic adaptations.

The neural basis for slow responses to environmental changes and need states is often conceptualized in terms of a labeled-line organization, wherein distinct hypothalamic cell types play unique roles integrating external and internal sensation to drive adaptive responses specific to each need (Joëls & Baram, 2009; Ulrich-Lai & Herman, 2009; Sternson, 2013; Saper & Lowell, 2014; Herman & Tasker, 2016; Biran et al., 2018). In contrast, less is known about the neural basis of fast avoidance responses to the onset of environmental stressors. A role for the hypothalamus in these fast behaviors is not well established; for instance, it could be the case that acute stressors directly recruit locomotor escape circuits through external or internal sensory pathways that bypass the hypothalamus altogether. If the hypothalamus is involved, it is unknown whether distinct cell types are acutely driven by each type of stressor in a labeled-line organization, or whether the neural encoding of acute threats is organized differently. This question has been challenging to investigate in mammals, because cellular-resolution functional recordings of hypothalamic neurons are rare; when achieved, these recordings are often limited to unclassified cell types or a single molecularly-defined cell type at a time (Lin et al., 2011; Jennings et al., 2015; Remedios et al., 2017; Zimmerman et al., 2019). Furthermore, deep-brain recordings in rodents typically involve invasive procedures that destroy a large amount of brain tissue in order to gain electrical or optical access to the hypothalamus.

To address these challenges, we chose to investigate fast-timescale encoding of environmental stressors in the larval zebrafish. Adult and larval zebrafish demonstrate both slow endocrine adaptations and fast behavioral avoidance in response to environmental stressors, including heat, salinity, and acidity, among others (Wendelaar Bonga, 1997; Hoshijima & Hirose, 2007; Yeh et al., 2013; De Marco et al., 2014; Kwong et al., 2014; Schulte, 2014; Haesemeyer et al., 2015; 2018; Schreck & Tort, 2016; vom Berg-Maurer et al., 2016; Takei & Hwang, 2017; Wee et al., 2019). The structure and function of the hypothalamus is conserved across fish and mammals (Chiu & Prober, 2014; Biran et al., 2015), and larval zebrafish possess a number of experimental advantages over rodents. Notably, the activity of single neurons across the entire brain can be observed through Ca^2+^ imaging without invasive surgery, to determine the location of neurons with activity correlated to behavioral events (Ahrens et al., 2012; Ahrens & Engert, 2015). Furthermore, we can apply our recently established methods for linking cellular-resolution neural activity during behavior to the molecular identity of the exact same cells in the same animal (Lovett-Barron et al., 2017). Finally, the environment around these small aquatic animals can be changed in controlled settings with high temporal resolution, to levels approximating the extremes recorded in their natural environments (Parichy, 2015). Together, these features allow for testing the correspondence between molecularly-defined cellular identity and the physiological and behavioral roles of the same neurons in the fast avoidance response to diverse and distinct environmental stressors.

We observed that larval zebrafish execute a common avoidance-like turning response at the onset of transient environmental stressors, including rapid increases in heat, salinity, or acidity. Brain-wide cellular-resolution Ca^2+^ imaging revealed distributed representations of stressor detection and resultant motor action, with notable stimulus-encoding in the preoptic hypothalamus – a neurosecretory region homologous to the paraventricular hypothalamus in mammals (Herget et al., 2014; Biran et al., 2015; 2018) and well-known to be involved in slow homeostatic adaptation (Joëls & Baram, 2009; Ulrich-Lai & Herman, 2009; Herman & Tasker, 2016). To explore the neuronal dynamics of molecularly-defined cell types within the hypothalamus during this behavior, we extended a method we had previously developed (MultiMAP; Lovett-Barron et al., 2017) to enable the cellular-resolution alignment of neural activity imaging to multiplexed gene expression. We developed and applied a next-generation form of this method to record from six to nine cell types at once, defined by the expression of genes for neuropeptide transmitters.

We found that heat, salinity, and acidity drove distinct population dynamics in the preoptic hypothalamus. However, the responses to each stressor were not confined to a single classical cell type, but instead were distributed across multiple peptidergic cell types. Despite their many differences, we found that classically distinct peptidergic neuron types shared several common features: their activation promoted avoidance behavior, they co-expressed glutamatergic genes, and their outputs converged upon a set of brainstem spinal-projecting neurons whose glutamatergic inputs were required for avoidance of environmental stressors. Ablation of each individual peptidergic cell type did not disrupt the fast avoidance behavior, whereas broader ablation of the peptidergic neurons in the region did reduce avoidance.

Together, these results reveal that multiple distinct types of peptidergic neurons in the hypothalamus are capable of generating a common rapid avoidance response at the onset of diverse environmental stressors, through shared convergence in the brainstem. This fast, cell-type invariant function complements these cells’ well-known capacity to generate slower, specific homeostatic adaptations to chronic stress. The identity of functional “cell types” in this context is therefore not static, but instead a flexible categorization of neuronal activity and behavioral contributions that can change across different timescales.

## Results

### Rapid, pituitary-independent avoidance of multiple stressors

We exposed larval zebrafish to environmental stressors that threaten homeostasis using transient (40 s) increases in water salinity (+50 mM NaCl), acidity (+0.1 mM HCl, to pH 4.8), or heat (+7 °C); fish were partially restrained in agarose, so we were able to monitor motor behavior in a configuration compatible with functional neural imaging (Figure 1a, Figure S1a, Methods). Each stressor increased the rate of turning behavior within 20 seconds of stimulus onset (Figure 1b; salinity 1.91±0.28 movements, acidity 1.47±0.31 movements, heat 1.55±0.28 movements, blank/no stimulus 0.58±0.24 movements, mean±s.e.m., N=57 fish), but not forward swimming (Figure S1b; salinity 0.28±0.10 movements, acidity 0.25±0.11 movements, heat 0.26±0.11 movements, blank/no stimulus 0.25±0.10 movements, mean±s.e.m., N=57 fish), consistent with avoidance responses observed in freely-moving animals (De Marco et al., 2014; vom Berg-Maurer et al., 2016). These responses did not habituate over multiple trials (5 trials per fish, Figure S1c; p> 0.1, Wilcoxon signed-rank tests). The similarity between behavioral actions evoked by different stressors suggested that the zebrafish were executing a common or generic avoidance response.

**Figure 1.**
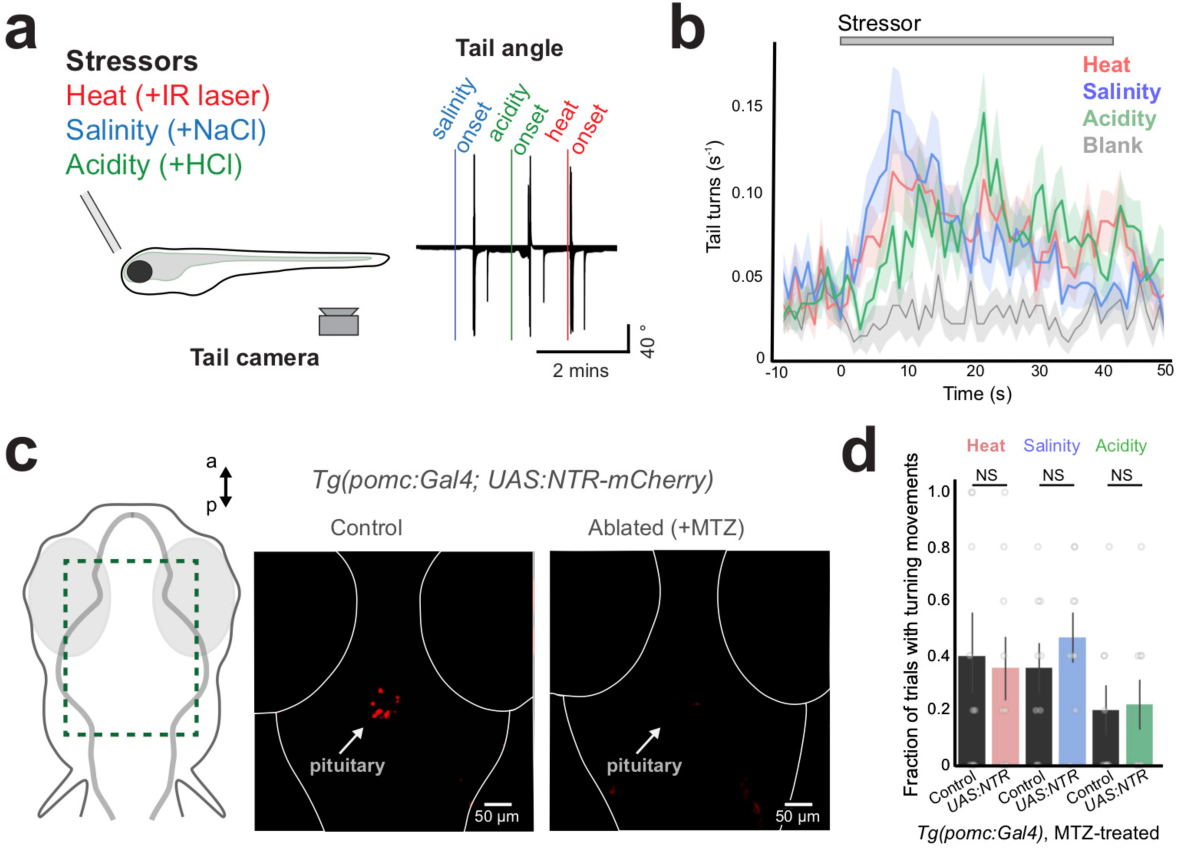
Rapid, pituitary-independent avoidance response to the onset of environmental stressors. **a)** Schematic of experiment (left). Head-restrained larval zebrafish are presented with 40 s stressors (30 s no-stimulus interval), while tail movements are recorded. Example tail angle of a fish during stressor presentation, showing turn-or escape-like movements to each stressor type (right). **b)** Rate of turning movements evoked by each stressor and a blank stimulus (no change in environment). Mean ± SEM, 1 s bins, N = 57 fish (average of 5 trials each). **c)** Ventral sections of *Tg(pomc:Gal4;UAS:NTR-mCherry)* fish, showing ablation of *pomc*^+^ cells in the pituitary (red) upon treatment with metronidazole (MTZ; right). **d)** Summary data from control and ablated fish. N = 9 fish per group, mean ± SEM, individual fish are points. Mann-Whitney U tests, corrected for multiple comparisons. NS = “not significant”, all p > 0.4.

Rapid avoidance responses are generally hypothesized to be independent of slower endocrine adaptations, which are driven by the systemic release of cortisol and other stress hormones into the circulatory system via the pituitary (Joëls & Baram, 2009; Ulrich-Lai & Herman, 2009; Herman & Tasker, 2016; Biran et al., 2018). In agreement with this hypothesis, we found that ablation of corticotrophs in the anterior pituitary (Liu et al., 2003; Davison et al., 2007) (Figure 1c, Figure S1d, Methods), a necessary component of the hormonal stress response, did not disrupt fast behavioral responses to heat, salinity, or acidity (Figure 1d, Figure S1e; all p>0.4, Mann-Whitney U tests corrected for multiple comparisons). These data indicate that zebrafish execute a rapid, common behavioral avoidance response to the onset of environmental stressors through a pituitary-independent pathway, prompting us to search for the central brain regions involved in this fast avoidance behavior.

### Brain-wide imaging of stressor-evoked neural dynamics

Slow homeostatic adaptations to environmental stressors are known to engage a distributed network in vertebrate brains, including the hypothalamus and various areas of the forebrain and brainstem (McEwen, 2007; Joëls & Baram, 2009; Ulrich-Lai & Herman, 2009; Herman & Tasker, 2016). To determine which brain regions are involved in rapid stressor-avoidance responses in zebrafish, we performed brain-wide two-photon Ca^2+^ imaging in fish with pan-neuronal expression of GCaMP6s (*Tg(elavl3:H2B-GCaMP6s)*, Vladimirov et al., 2014) as they responded to the onset of environmental stressors (Figure 2a,b, Figure S2a, Methods). To provide an avoidance-promoting stimulus independent of homeostatic stress, we also included a visual looming stimulus (Temizer et al., 2015; Dunn et al., 2016; Lovett-Barron et al., 2017).

**Figure 2.**
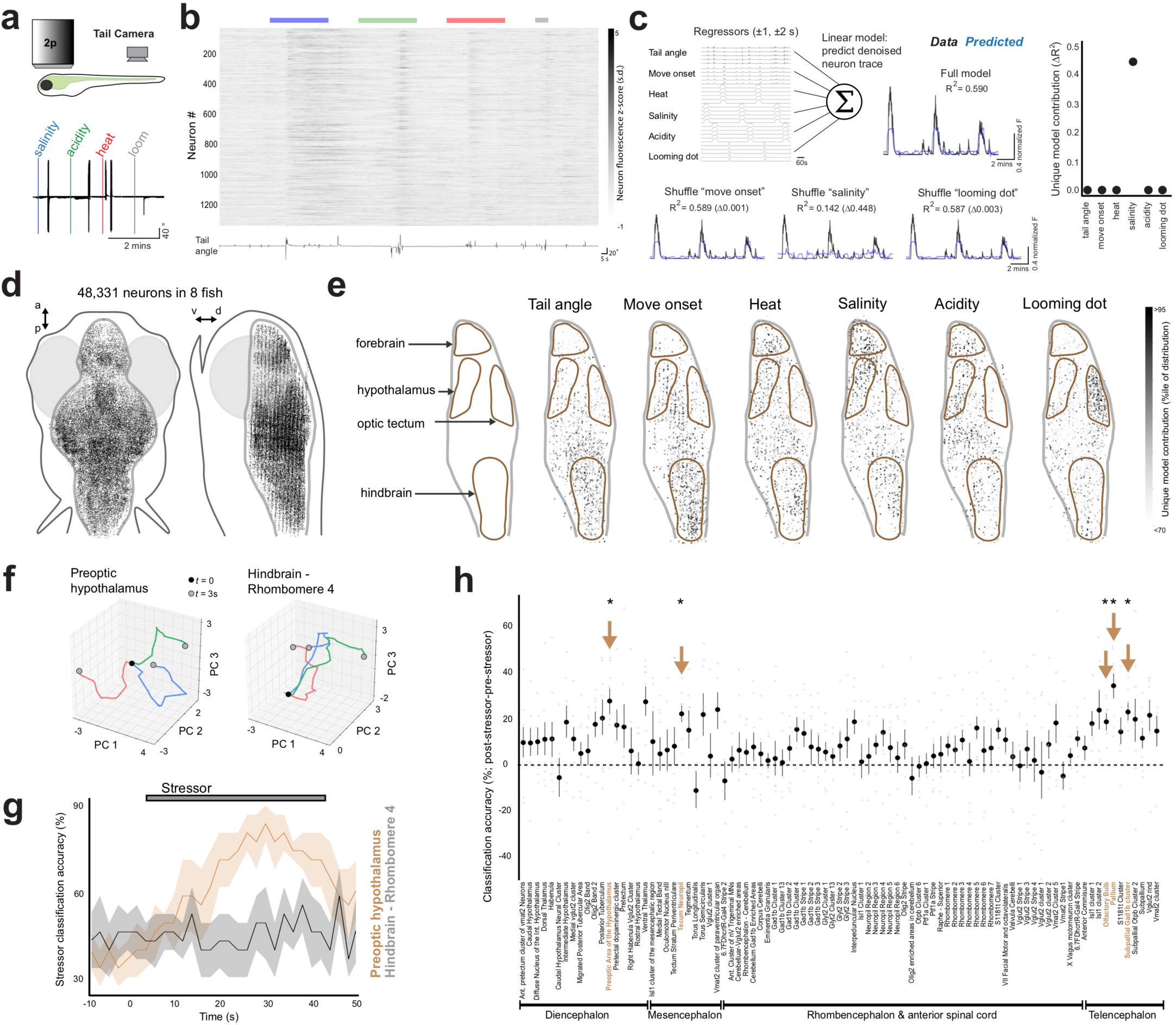
Brain-wide imaging reveals rapid encoding of stressors by neural populations in the hypothalamus and forebrain. **a)** Schematic of experiment, and example tail angle over stressor presentation. **b)** All neurons recorded with two-photon Ca^2+^ imaging in an example *Tg(elavl3:H2B-GCaMP6s)* fish, with the times of stressors (above) and tail angle (below) for an example trial (5 such trials during imaging experiment). **c)** Schematic of single-neuron analyses, and data from an example neuron. A linear model of each cell’s denoised fluorescence time series is composed from behavioral events (convolved with calcium indicator decay), and the importance of each event is determined by the change in model performance (ΔR^2^) upon shuffling of each event time-series (unique model contributions, UMC). UMC plot for the example neuron (right). This example neuron preferentially encodes salinity. **d)** Top and side projections show location of analyzed neurons from all fish (N = 48,331 cells from 8 fish), registered to a common brain volume. **e)** Spatial distribution of UMC for each variable, shown in side projection of the medial 40 µm of volume. Values are displayed as the percentile within the distribution of UMC for that task component across all cells. Brain regions are outlined in brown. **f)** Trajectory of population activity in two anatomical subregions of an example fish. Z-scored and baseline-subtracted time-series of the first three principal components is shown from stressor onset (black dot) to three seconds later (grey dots) (heat=red, salinity=blue, acidity=green). **g)** Accuracy of stressor classifier on held out data, derived from the top ten principal components in two anatomical regions (N = 8 fish, 2 s bins, mean ± SEM). **h)** Difference in mean classifier accuracy between the time after stressor onset compared to before stressor onset, across anatomical sub-regions (N = 8 fish). Arrowheads denote regions that are significantly different from zero (one-sample t-tests, corrected for multiple comparisons across all brain regions). *p<0.05.

To classify the responses of each recorded neuron, we constructed linear models based on time series of sensory stimuli and locomotor actions, and fit these models to the denoised activity traces from each neuron (Figure 2c, N = 48,331 neurons in 8 fish, Methods). Comparing the fit of complete models (R^2^) with the fit of partial models constructed from shuffling each task component produced an array of unique model contributions (UMC, ΔR^2^); each UMC value indicated the relative importance of that task component for modulating the activity of each neuron (Figure 2c) (Musall et al., 2018; Engelhard et al., 2019). To visualize the spatial distribution of UMC values for task-encoding neurons across the brain, we registered each brain-wide recording to a common atlas (Figure 2d,e, Figure S2b, Methods). This approach reproduced patterns of activity documented in previous studies, including midbrain and hindbrain neurons responsive to swimming movements (Ahrens et al., 2012; Vladimirov et al., 2014) and neurons in the optic tectum responsive to looming stimuli (Temizer et al., 2015; Dunn et al., 2016). Furthermore, we found salinity-, acidity, and heat-encoding neurons throughout the brain, with localization to similar anatomical regions, particularly the forebrain and hypothalamus (Figure 2e) – two areas classically associated with slower endocrine and behavioral adaptation to stressors and environmental perturbation (McEwen, 2007; Joëls & Baram, 2009; Ulrich-Lai & Herman, 2009; Herman & Tasker, 2016).

The overlap between brain areas recruited by heat, salinity, and acidity and the similarity between behavioral responses to each of these stressors (Figure 1b) prompted us to search for neural populations that distinguished between different stressors. To separately analyze the activity of populations within each brain region, we sorted neurons by anatomical location (Figure S2c) (Randlett et al., 2015) and used principal component analysis to visualize a low-dimensional representation of population activity for different brain regions (Figure 2f, Methods). To quantify which brain areas could distinguish among the types of stressors, we sought to classify the type of stimulus delivered on each trial (heat, salinity, acidity, or looming dot) based on regional population dynamics. Using the time-series of the top ten principal components in each region, we trained a classifier to distinguish stimulus type in bins of two seconds (Figure 2g, Methods) and quantified stimulus-driven classification across brain regions by the difference in post-stimulus versus pre-stimulus classification accuracy. We found that a small number of brain regions distinguished among these stimuli, including sensory regions associated with the stimuli used in this behavior (olfactory bulb and the retinal-recipient regions of the optic tectum), as well as regions of the pallium/subpallium and preoptic hypothalamus (Figure 2h, p<0.05, One-sample t-tests, corrected for multiple comparisons across all brain regions).

These data reveal that the onset of different environmental stressors is largely encoded by populations in overlapping regions of the hypothalamus and forebrain, and that their local population dynamics can rapidly distinguish among the types of stressors. The preoptic hypothalamus is of particular interest, since this is a neurosecretory region homologous to the paraventricular hypothalamus in mammals (Herget et al., 2014; Biran et al., 2015; 2018), plays an important role in slower adaption to homeostatic or environmental stressors (Joëls & Baram, 2009; Ulrich-Lai & Herman, 2009; Herman & Tasker, 2016), and exhibited the highest stressor classification accuracy across diencephalic structures (Figure 2h). Guided by the results of our brain-wide screen, we therefore focused subsequent efforts on studying this ancestral brain region in detail.

### Cellular registration of neural dynamics to gene expression

The neurosecretory preoptic hypothalamus (paraventricular nucleus in mammals) is a structure rich in peptidergic neurons, including cells that secrete oxytocin, vasopressin, and/or corticotrophin-releasing factor, among several other neuropeptides (Joëls & Baram, 2009; Herget & Ryu, 2015; Romanov et al., 2017). We initially hypothesized that distinct cell types may selectively respond to distinct types of stressors, as has been suggested for the slower neuroendocrine adaptation to environmental and homeostatic stressors (Joëls & Baram, 2009; Ulrich-Lai & Herman, 2009; Sternson, 2013; Saper & Lowell, 2014; Herman & Tasker, 2016). To test this hypothesis, we sought to record from multiple identified peptidergic neurons at once, from head-restrained zebrafish behaviorally responding to transient environmental stressors. We focused on cell types defined by the expression of key neuropeptide transmitters common across both the zebrafish and mammalian hypothalamus: oxytocin/isotocin (*oxt),* arginine vasopressin/vasotocin (*avp*), corticotrophin-releasing factor (*crf*), neuropeptide-Y (*npy)*, vasoactive intestinal polypeptide (*vip*), and somatostatin (*sst*).

In order to record from multiple molecularly defined neurons at once, we expanded our MultiMAP approach (Lovett-Barron et al., 2017) to perform cellular-resolution registration of neural activity imaging to *in situ* multiplexed gene expression (Figure 3a). The experimental protocol can be summarized as follows: we first performed live volumetric neural activity imaging in the preoptic hypothalamus of *Tg(elavl3:H2B-GCaMP6s)* fish during behavior (Figure 3b), after which fish were preserved in a fixative. We next performed multiple rounds of intact-tissue molecular phenotyping using whole-animal triple fluorescent *in situ* hybridization (hybridization chain reaction, HCR; Choi et al., 2018, Supplementary Table 1). After each round of fluorescent *in situ* hybridization, we imaged the preoptic hypothalamus again to collect volumes of GCaMP neurons as well as *in situ* hybridization labels. We then digested these DNA-based labels and repeated this procedure (Figure 3a). Our method ensured preservation of GCaMP6 fluorescence throughout the brain after fixation, hybridization, label digestion, and re-labeling (Methods). This stable GCaMP signal, the common cellular label across imaging iterations, allowed us to use a non-rigid registration approach (Rohlfing & Maurer, 2003; Lovett-Barron et al., 2017) to align the fixed brain volumes (and associated *in situ* hybridization labels) with the live brain volume at cellular resolution.

**Figure 3.**
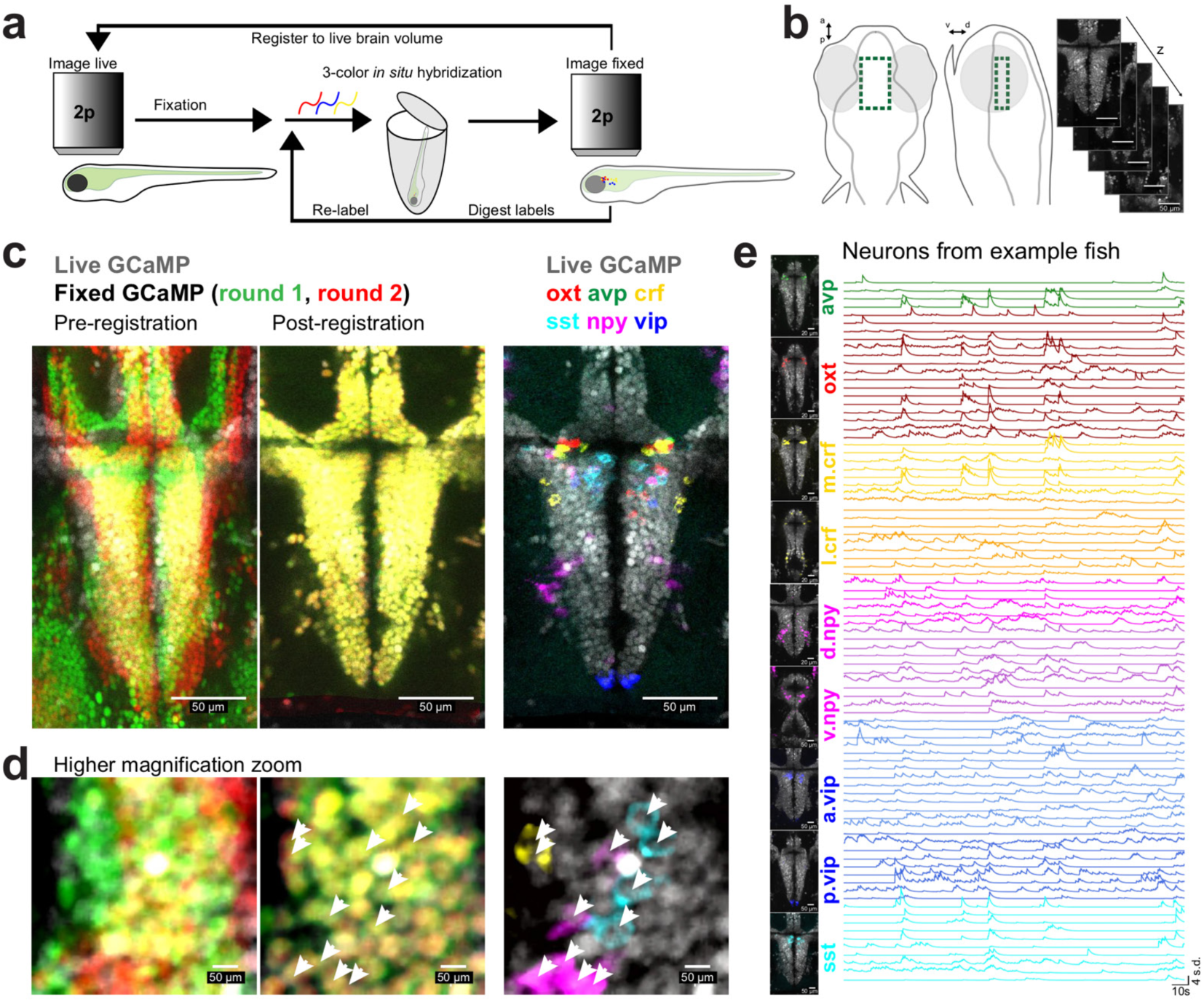
Simultaneous recording of multiple peptidergic cell types in the hypothalamus through cellular-resolution registration of multiplexed gene expression and neural dynamics. **a)** Schematic of process for registration of multiple rounds of multicolor fluorescent *in situ* hybridization (fISH) to cellular-resolution live-brain imaging during behavior. **b)** Location of volumetric two-photon functional imaging encompassing the preoptic hypothalamus (top and side views), and functional imaging z-planes from an example *Tg(elavl3:H2B-GCaMP6s)* fish. **c)** Select z-plane from volumes in an example fish, showing live functional imaging (grey GCaMP signal) overlapping with GCaMP signal after fixation and both rounds of *in situ* hybridization (left: pre-registration, right: post-registration) or with *in situ* hybridization markers of neuropeptides from both rounds (right). **d)** Same fish as shown in panel c, but higher magnification of a small area. White arrowheads indicate labeled cells. **e)** Location of analyzed neurons and fluorescence activity traces of simultaneously recorded labeled neurons in an example fish. *Crf* is split into medial (*m.npy*) and lateral (*l.npy*) populations. *Npy* is split into dorsal (*d.npy*) and ventral (*v.npy*) populations. *Vip* is split into anterior (*a.vip*) and posterior (*p.vip*) populations.

These methods allowed us to overlay six to nine *in situ* hybridization labels onto the live brain volume for each fish studied (Figure 3a,c,d, Figure S3a,b), whereupon we could classify recorded neurons by their molecular identity and anatomical location within the preoptic hypothalamus. By retrospectively identifying the activity time-series for each classified peptidergic neuron, we were able to analyze the dynamics of many molecularly-defined neurons that had been recorded simultaneously during behavior (Figure 3e). Confirming previous studies in the larval zebrafish (Herget & Ryu, 2014), we found many co-labeled *avp*^+^/*crf*^+^ neurons around the third ventricle (Figure 3c, Figure S3c, 36.4% of *avp*^+^ cells are also *crf*^+^).

We were able to register at least three rounds of triple fluorescent *in situ* hybridization to the live brain volume, amounting to simultaneous activity imaging of cell types defined by the expression of nine distinct genes (Figure S4). We designed these methods for compatibility with further rounds of hybridization, imaging, and digestion – thus potentially allowing for future studies using barcoded *in situ* sequencing or *in situ* hybridization approaches such as STARmap (Wang et al., 2018) or MERFISH (Moffitt et al., 2018). This general strategy may thus enable the merging of large-scale neural activity recording during behavior with expression profiling of large numbers of genes.

### Environmental stressors engage multiple peptidergic cell types

We analyzed the activity of individual molecularly-defined neurons using the same linear models used for brain-wide recordings – establishing encoding preference of each neuron based on the unique model contributions of each task component (UMC, ΔR^2^). We then clustered neurons by these UMC arrays (Methods), to examine the molecular identity of functionally similar neurons encoding heat, salinity, acidity, looming dots, and/or movement parameters (Figure 4a). In contrast to our initial hypothesis that each stressor would be narrowly encoded by a unique set of peptidergic cell types, we observed that each stressor recruited neurons across molecularly-defined cell classes. As a consequence, each peptidergic cell type was composed of neurons that were preferentially active during diverse sensory stimuli and/or motor actions (Figure 4b).

**Figure 4.**
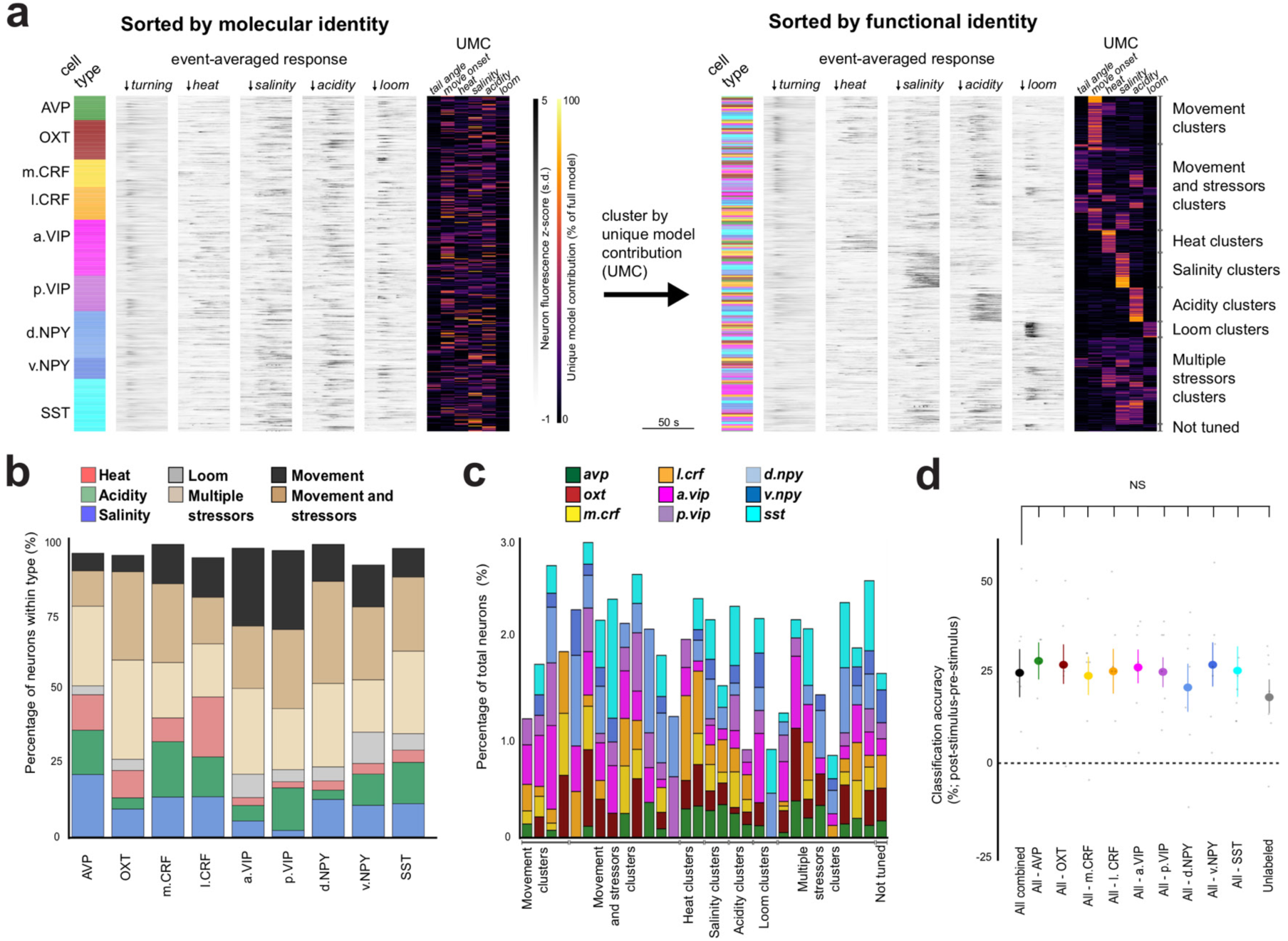
Distributed encoding of stressors and motor action across multiple hypothalamic cell types. All identified peptidergic neurons recorded (N = 452 peptidergic neurons from 28,211 total recorded cells in 7 fish), with cell type denoted in color, event onset-averaged responses as grey intensity, and unique model contributions (UMC) values from linear model fit displayed as a heatmap (% of full model). Left: neurons sorted by molecular and anatomical identity. Right: after clustering UMC values, same neurons sorted by functional identity (cluster ID, written at right). **b)** Cluster identities for each molecular/anatomical cell type examined (N = 7 fish, number of neurons in each cell type = 33, 53, 37, 44, 75, 48, 63, 28, and 71, from left to right). “Movement” encompasses “tail angle” and “move onset” clusters. The “not tuned” cluster is not displayed. **c)** The cell types within each functional cluster, plotted as percentage of all neurons in the cluster (including unlabeled neurons). **d)** Difference in mean stressor classifier accuracy between the time after stressor onset compared to before stressor onset, across cell types (N = 7 fish). Data shown for all peptidergic neurons together, and with neurons from individual cell classes removed. Mean ± SEM, individual fish are points. All comparisons are paired t-tests with the “All combined” group (all recorded peptidergic neurons), corrected for multiple comparisons. NS = “not significant”, all p > 0.3.

While we observed a diversity of functional response types within every molecularly-defined cell type, we also noted that not every cell type displayed the same distribution of preferences at the single neuron level. For example, we observed that a higher percentage of *avp*^+^ and *crf*^+^ neurons were purely responsive to heat, salt, or acidity (48.5% of *avp*^+^ neurons, 40.5% of medial *crf*^+^ neurons, and 47.7% of lateral *crf*^+^ neurons), many *npy*^+^ neurons were purely movement-responsive (26.7% of dorsal *npy*^+^ neurons, and 27.1% of ventral *npy*^+^ neurons), and most *oxt, vip*, and *sst* neurons showed mixed selectivity (67.9% of *oxt*^+^ neurons, 68.3% of anterior *vip*^+^ neurons, and 59.2% of *sst*^+^ neurons) (Figure 4b). In general, while multiple cell types were found in each functional cluster, we observed heterogeneity in the exact set of peptidergic neurons in each (Figure 4c). These results appear consistent with previous reports of hypothalamic imaging of single cell types in zebrafish, where *oxt*^+^ neurons were found to be responsive to noxious stimuli and/or motor responses (Wee et al., 2019), and a subset of *crf*^+^ neurons responded to salinity and acidity (vom Berg-Maurer et al., 2016). Both of these studies focused their examinations on a single molecularly-defined cell type, but noted broad recruitment of unlabeled neurons. By recording from many molecularly-defined cell types at once, here we show that neural encoding of diverse stressors and the resultant motor responses is distributed across a number of distinct peptidergic neuron types in the preoptic hypothalamus.

Despite the differences we observed across cell types, the diversity of functional responses within each cell type suggested that none of these peptidergic neurons plays a unique role in the classification of environmental stressors. We tested this interpretation by asking whether the absence of each cell type individually would reduce the capacity for the overall peptidergic neuron population to classify stimulus type (Methods). For each fish, we used the time-series of all peptidergic neurons to train a classifier to distinguish stressor type, then repeated these analyses with each peptidergic cell type removed from the dataset. We found that individually removing any of the peptidergic cell types indeed was insufficient to reduce the classification accuracy of the overall population (Figure 4d; p>0.3, paired t-tests with the “All combined” group (all recorded peptidergic neurons), corrected for multiple comparisons).

These experiments indicate that during the onset of environmental stressors, cell types in the neurosecretory hypothalamus do not show a strict correspondence between neural activity patterns and molecular identity, at least with respect to neuropeptide gene expression. While we did find subtle differences among cell types, each molecularly-defined cell type contained neurons with diverse functional properties, and none of these cell types played a unique role in the hypothalamic population response to each specific environmental stressor.

### Cell type manipulations reveal overlapping roles

Our functional imaging data revealed that molecularly-distinct hypothalamic cell types exhibited similar neural activity patterns during behavior. This observation raised the possibility that these different cell types may play similar roles in the behavioral avoidance response to the onset of environmental stressors. We tested this idea by manipulating these cell types in behaving animals, using the Gal4/UAS system (Kawakami et al., 2016) to drive transgene expression in restricted subsets of peptidergic neurons in the preoptic hypothalamus (Figure 5a,b, Figure S5a, Methods). We used three different transgenic lines that labeled neurons according to the expression of a specific neuropeptide: oxytocin/isotocin (*Tg(oxt:Gal4)*, Wee et al., 2019), corticotrophin-releasing factor (*Tg(crf:Gal4)*, Methods), and somatostatin (*Tg(sst3:Gal4)*, Förster et al., 2017), as well as a line that labels multiple peptidergic neurons (*Tg(otpba:Gal4)*, Fujimoto et al., 2011, Herget & Ryu, 2015). We first asked whether artificial stimulation of these cell types would induce avoidance-like motor actions in the absence of a stressor. We used optogenetics to activate these neurons (Yizhar et al., 2011), directing a fiber optic to the preoptic hypothalamus (Figure 5c, Figure S5b) and activating channelrhodopsin (ChR2)-expressing neurons with blue light at 5 Hz (Methods). We observed dose-dependent light-evoked tail turning when we stimulated *Tg(oxt:Gal4; UAS:ChR2-mCherry), Tg(crf:Gal4; UAS:ChR2-mCherry)*, or *Tg(otpba:Gal4; UAS:ChR2-mCherry)* fish, but not *Tg(sst3:Gal4; UAS:ChR2-mCherry)* fish (Figure 5d; 0.5 mW: p > 0.2 for all lines, 6 mW: p < 0.05 for *Tg(otpba:Gal4)*, 14 mW: p < 0.05 for *Tg(oxt:Gal4), Tg(crf:Gal4*), and *Tg(otpba:Gal4)*, Mann-Whitney U tests, corrected for multiple comparisons). Therefore, specific stimulation of multiple different hypothalamic cell types suffices to induce behavioral output similar to that observed during the onset of environmental stressors that naturally activate those same neurons.

**Figure 5.**
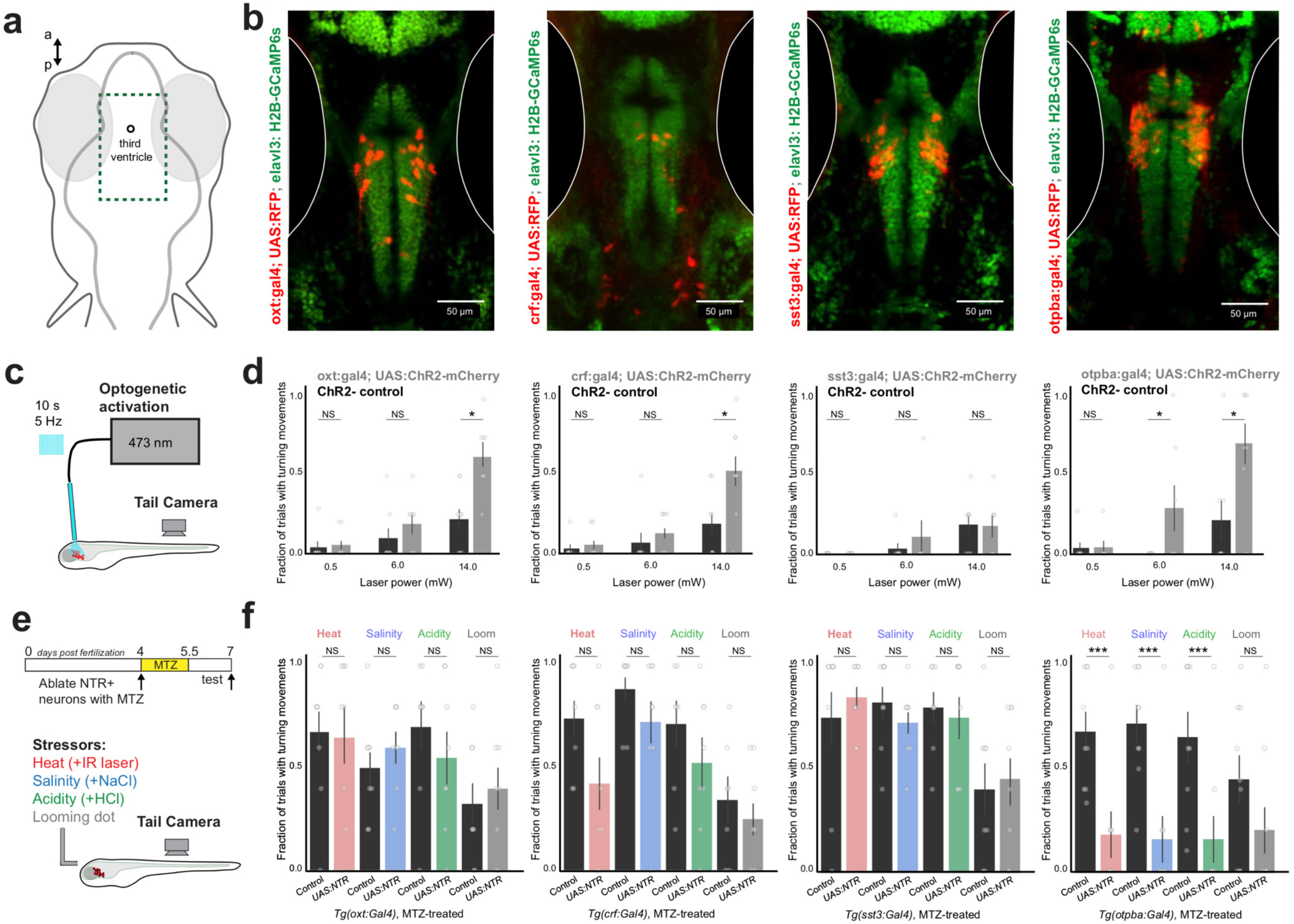
Overlapping functional roles for multiple hypothalamic peptidergic cell types. **a)** Location of images in panel b - encompassing the preoptic hypothalamus. **b)** Location of Gal4^+^ neurons in transgenic lines used to target peptidergic cell classes. Gal4 drives “UAS:RFP” (UAS:NTR-mCherry or UAS:ChR2-mCherry), in a background of *Tg(elavl3:H2B-GCaMP6s)*. **c)** Schematic of experiment - measuring the behavioral response to focal optogenetic activation of the preoptic hypothalamus, in the absence of other stimuli. **d)** Results of optogenetic activation in ChR2+ and ChR2-fish, for each Gal4 line. Mean ± SEM, individual fish are points. N = (8,8), (7,8), (8,8), (7,8) – (ChR2+, ChR2-), left to right. Mann-Whitney U tests, corrected for multiple comparisons within genotype. NS = “not significant”, all p > 0.2. *p < 0.05. **e)** Schematic and timeline of experiment - using 36 hours of 10 mM metronidazole (MTZ) treatment to ablate NTR+ neurons, before testing behavior in response to stressors. **f)** Results of behavioral experiments in NTR+ and NTR-fish, for each transgenic line, all treated with MTZ. Mean ± SEM, individual fish are points. N = (8,8), (8,7), (8,8), (9,9) – (NTR+, NTR-), left to right. Mann-Whitney U tests, corrected for multiple comparisons within genotype. NS = “not significant”, all p > 0.1. ***p<0.005.

We next tested whether these cell types would be required for the natural avoidance-like turning response to environmental stressors. Using a chemogenetic approach to ablate nitroreductase (NTR)-expressing neurons with metronidazole (Figure 5e, Figure S5c; Methods; Davison et al., 2007), we found that behavioral responses were unchanged after ablation of each individual cell type in *Tg(oxt:Gal4; UAS:NTR-mCherry), Tg(crf:Gal4; UAS:NTR-mCherry)*, or *Tg(sst3:Gal4; UAS:NTR-mCherry)* fish (Figure 5f; all p>0.1, Mann-Whitney U tests, corrected for multiple comparisons). However, we found that ablation of neurons in the broader-expressing *Tg(otpba:Gal4; UAS:NTR-mCherry)* line reduced the response to heat, salinity, and acidity (all p<0.005, Mann-Whitney U tests, corrected for multiple comparisons), but not looming stimuli (p>0.1) (Figure 5f). In all conditions, ablation did not disrupt the ability for fish to physically execute large-angle tail movements (Figure S5d; all p>0.1, Mann-Whitney U tests, corrected for multiple comparisons).

In addition to labeling peptidergic neurons in the neurosecretory preoptic hypothalamus, the *Tg(otpba:Gal4)* line also labels dopaminergic neurons in the posterior tuberculum (Löhr et al., 2009). However, ablation of these neurons in *Tg(th:Gal4; UAS:NTR-mCherry)* fish (Li et al., 2015) did not disrupt behavior relative to controls (Figure S5e; all p>0.4, Mann-Whitney U tests, corrected for multiple comparisons). In contrast, restricted optogenetic inhibition of preoptic *otpba*^+^ neurons using the inhibitory channelrhodopsin GtACR1 (Mohamed et al., 2017) in *Tg(optba:Gal4; UAS:GtACR1-eYFP)* fish (Methods) did reduce the behavioral response to environmental stressors (Figure S5f; p<0.05 for acidity, salinity, and heat, p>0.1 for loom, Mann-Whitney U tests, corrected for multiple comparisons). Together, this data suggests that *otpba*^+^ neurons in the preoptic hypothalamus, which encompasses *oxt*^+^, *avp*^+^, and *crf*^+^ neurons (Figure S5a; Herget et al., 2014) are required for avoidance responses to environmental stressors. We do not rule out the possibility, however, that other types of peptidergic neurons co-labeled by the *otpba* expression pattern also contribute to this behavior.

### Functionally similar cell types converge in the brainstem

The cell type-specific manipulation experiments revealed that different peptidergic cell types in the hypothalamus can play overlapping roles in the generation of rapid avoidance responses to environmental stressors. The similarity of behavioral effects we observed upon manipulation of different peptidergic neurons prompted us to search for morphological features shared by these cell types. We reasoned that hypothalamic cells may be capable of driving rapid locomotor behavior by activating premotor neurons that project to the spinal cord (Kimmel et al., 1982; Orger et al., 2008). To test this possibility, we labeled spinal projection neurons (SPNs) with injection of a red fluorescent dextran into the spinal cord of fish that express GFP in each Gal4 line (Figure 6a, Figure S6a, Methods). While axons from each set of neurons appears to ramify throughout the hindbrain region, we found that neurons labeled in *Tg(oxt:Gal4; UAS:GFP), Tg(crf:Gal4; UAS:GFP), Tg(sst3:Gal4; UAS:GFP)*, and *Tg(otpba:Gal4; UAS:GFP)* lines made prominent projections to a specific set of lateral spinal projection neurons known as RoL1 (*rostral lateral interneuron 1*, Figure 6b), which appose noradrenergic neurons of the locus coeruleus at their anterior/medial edge (Figure S6b).

**Figure 6.**
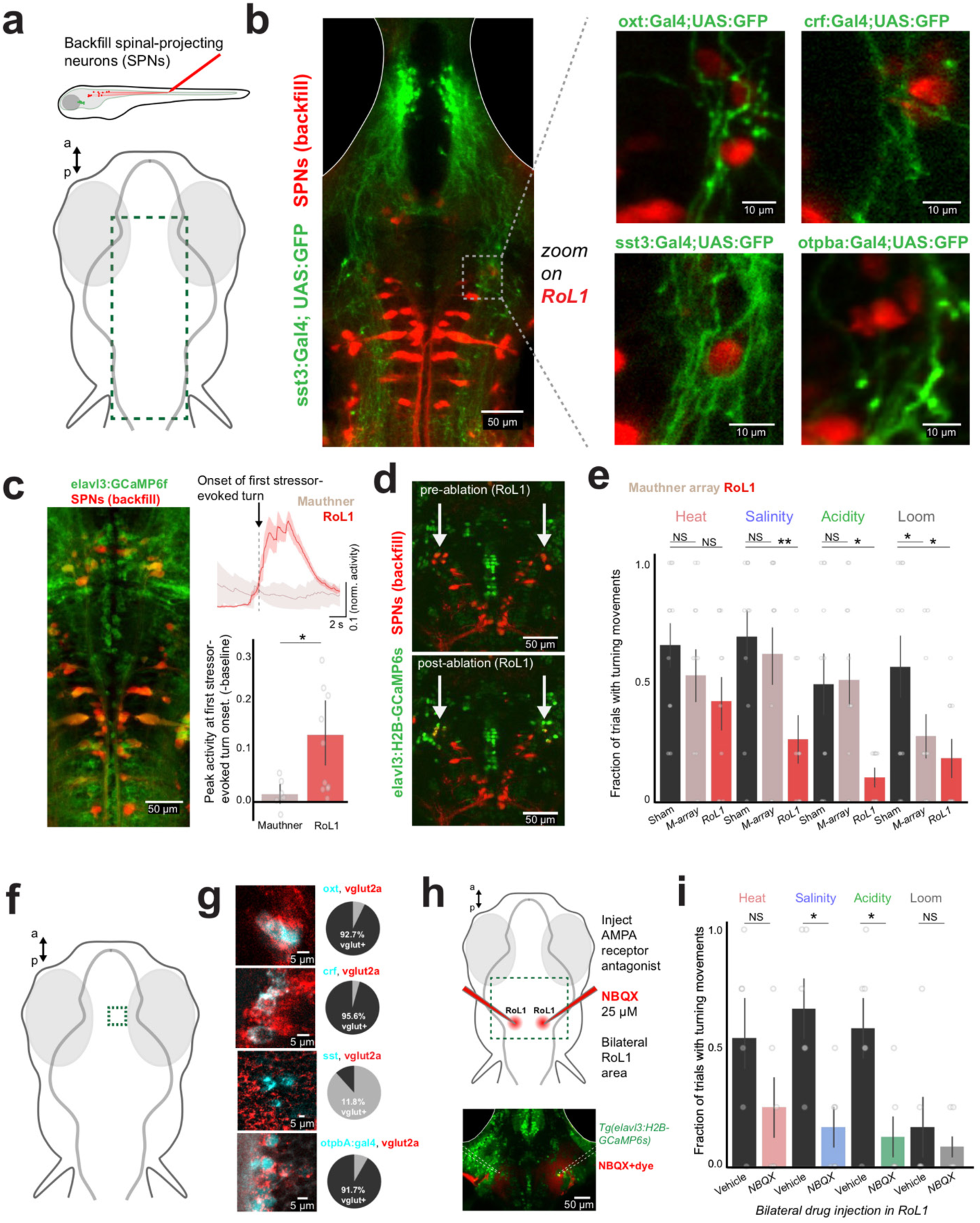
Functionally similar peptidergic cell types are glutamatergic and converge upon brainstem neurons required for avoidance. **a)** Schematic of spinal-projection neuron labeling, and location of image in panel b. Spinal-projection neurons (SPNs) were backfilled with Texas red dextran in fish expressing GFP in each cell type. Z-projection of example *Tg(sst3:Gal4; UAS:GFP)* fish (left). Overlap of GFP+ axons with back-labeled RoL1 neurons in each Gal4 line, with higher magnification of a small area (right). **c)** Two-photon imaging of SPNs in *Tg(elavl3:GCaMP6f)* fish. Z-projection of example fish (left) and mean activity of two SPN cell classes in example fish, aligned to the onset of the first turn evoked by salinity, acidity, or heat (top). Summary of peak activity in the 2 s around stimulus onset, normalized to mean fluorescence 2 s prior (bottom). Mean ± SEM, individual fish are points. Paired-sample t-test (N = 4 fish, N=7,11 cells). *p < 0.05. **d)** Example of two-photon ablation - before and after bilateral ablation of backfilled RoL1 neurons (noted by arrows). **e)** Results of behavioral experiments in ablated (RoL1 neurons or Mauthner-array (M-array) neurons) or sham-ablated fish. Mean ± SEM, individual fish are points. N = (11,11,10) – (sham, M-array, RoL1), left to right. Mann-Whitney U tests, corrected for multiple comparisons. NS = “not significant”, all p > 0.1. *p < 0.05, **p<0.01. **f)** Image location in panel g - within the preoptic hypothalamus. **g)** Multi-color fluorescent *in situ* hybridization shows overlap of neuropeptide expression (cyan) with *vglut2a* (red). At bottom is overlap of *vglut2a* (red) with GFP expression in *Tg(otpba:Gal4; UAS:GFP)* fish. Pie charts showing number of peptide^+^ neurons that are also *vglut2a*^+^ (4 sections from 2 fish each). **h)** Schematic of experiment, with location of NBQX injection around RoL1 (top). Image of drug spread in an example fish, visualized by red dye (bottom). **i)** Results of behavioral experiments in RoL1-injected fish (NBQX or vehicle). Mean ± SEM, individual fish are points. N = (6,6) – (vehicle, NBQX). Mann-Whitney U tests, corrected for multiple comparisons. NS = “not significant”, all p > 0.1. *p < 0.05.

RoL1 neurons have been previously observed to be active during escape behaviors (Gahtan et al., 2002), but activity during homeostatic stressor-evoked avoidance has not been documented. We therefore imaged the activity of spinal-projection neurons in *Tg(elavl3:GCaMP6f)* fish during behavior (Figure 6c). By aligning the activity of neurons with the first movement during heat, salinity, or acidity presentation, we found that RoL1 neurons were active at the onset of these homeostatic stressor-induced movements whereas the classical rapid escape neurons, the Mauthner cells (Korn & Faber, 2005), were not (Figure 6c; baseline-subtracted normalized response, RoL1 = 0.14±0.04, Mauthner = 0.02±0.01, mean±s.e.m.). To determine if RoL1 neurons are required for stressor-evoked avoidance responses, we selectively removed these neurons (∼3-5 neurons per hemisphere) using two-photon single-cell ablation (Figure 6d, Figure S6c, Methods). This manipulation reduced the behavioral response to salinity, acidity, and looming stimuli (p<0.05, Mann-Whitney U tests, corrected for multiple comparisons), while ablation of Mauthner neurons and their segmental homologues (Figure S6d) only reduced the response to looming stimuli (Figure 6e, Figure S6e; p<0.05, Mann-Whitney U test, corrected for multiple comparisons), as has been previously described (Dunn et al., 2016). These data are also consistent with recent work demonstrating that Mauthner cell ablation does not disrupt pain-evoked escapes (Wee et al., 2019).

We next searched for the mechanisms linking the hypothalamus and RoL1 activation. While hypothalamic neurons can vary greatly in their production of specialized neuropeptides, these cells can also be more coarsely defined by the co-expression of classical amino acid transmitters such as glutamate or GABA (Schöne & Burdakov, 2012; Romanov et al., 2017; Moffitt et al., 2018). Examining the preoptic hypothalamus around the third ventricle, we found that most *oxt*^+^ and *crf*^+^ neurons were also *vglut2a*^+^ (92.7% and 95.6% *vglut2a*^+^, respectively), suggesting that these neurons co-release glutamate, whereas the majority of *sst*^+^ neurons were not *vglut2a*^+^ (11.8% *vglut2a*^+^) (Figure 6f,g, Methods). We also observed that the majority of preoptic hypothalamus neurons labeled by the *Tg(otpba:Gal4)* line were glutamatergic (91.7% *vglut2a*^+^). Therefore, the cell types that promoted turning behavior upon optogenetic stimulation (Figure 5d) likely co-release glutamate. To determine if RoL1 neurons are driven by glutamatergic input, we tested the behavior of fish after locally applying an ionotropic glutamate receptor blocker bilaterally to the RoL1 region (Figure 6h, Figure S6f, Methods). We found this manipulation reduced the behavioral response to salinity and acidity (Figure 6i, Figure S6g; p<0.05, Mann-Whitney U test, corrected for multiple comparisons). The residual behavioral response to heat after RoL1 neuron ablation or glutamate receptor blockade may be due to alternative pathways supporting heat-induced movement, such as direct projections from the trigeminal ganglia to the posterior hindbrain (Haesemeyer et al., 2018).

In sum, these data demonstrate that different types of peptidergic neurons in the preoptic hypothalamus share glutamatergic properties and converge upon a common set of spinal projection neurons in the brainstem – the RoL1 cells – that are required for avoidance. This convergent hypothalamus-brainstem-spinal pathway provides a circuit mechanism by which hypothalamic neurons that are classically considered to be “neuroendocrine” can quickly and directly influence behavior, independent of peptide secretion in the pituitary.

## Discussion

When animals encounter an environmental stressor that challenges homeostasis, they can adapt with a fast seconds-timescale avoidance response or a slow minutes-to-hours-timescale homeostatic adaptation (Wingfield, 2003; Heinrichs & Koob, 2004; Dallman, 2005; Joëls & Baram, 2009). Here we show that fast responses recruit some of the same types of neurons used for slow responses, particularly the neurosecretory hypothalamus. Taking advantage of whole-brain activity imaging in behaving larval zebrafish, we were able to find that neuronal populations in the neurosecretory preoptic hypothalamus rapidly discriminate increases in heat, salinity, or acidity to drive a stereotyped fast avoidance behavior at the onset of each stressor. Using an improved implementation of our MultiMAP technique (Lovett-Barron et al., 2017), we were able to merge neural activity imaging in the hypothalamus with multiplexed gene expression at cellular resolution, to discover that there was limited correspondence between molecular identity (defined by neuropeptide gene expression) and neural activity patterns in response to environmental stressors. Analysis of a subset of these cell types with transgenic lines revealed some similarities across neurons releasing the neuropeptides oxytocin and corticotrophin-releasing factor: their optogenetic activation drives avoidance behavior, they are commonly glutamatergic, and they converge on a small set of spinal-projecting premotor neurons that are essential for avoidance behaviors. Whereas chemogenetic ablation of each peptidergic cell type individually had no effect, broader peptidergic neuron loss-of-function in the region reduced avoidance responses to each stressor. Our results demonstrate that multiple distinct types of peptidergic neurons in the neurosecretory hypothalamus are capable of generating rapid avoidance responses to the onset of environmental stressors – a function complementary to these cells’ prominent role in slow homeostatic adaptation. Therefore, while specific neuropeptide expression may be the relevant dimension of gene expression to define functional cell types on slow timescales, we find that broader amino acid transmitter expression is a more relevant dimension to define functional cell types on faster timescales. These findings emphasize the importance of classifying functional units of the brain through an integrated description of individual cells based on gene expression, connectivity, and neural activity across a variety of timescales and behavioral conditions.

Key features of this system are consistent with studies of the mammalian hypothalamus. First, neurosecretory hypothalamic neurons have been reported to rapidly respond to a variety of stressful and aversive events, including oxytocin and corticotrophin-releasing factor neurons in the paraventricular nucleus (Condés-Lara et al., 2009; Kim et al., 2019), and vasopressin neurons in the supraoptic nucleus (Zimmerman et al., 2019). In addition, electrical stimulation of the paraventricular hypothalamus in mouse evokes rapid escape behaviors (Lammers at el., 1988), and optogenetic activation of corticotrophin-releasing factor neurons drives rodents to perform context-specific motor actions (Füzesi et al., 2016). Second, peptidergic neurosecretory neurons in the rodent paraventricular and supraoptic hypothalamus are known to co-express vesicular glutamate transporters, including the oxytocin, vasopressin, and corticotrophin-releasing factor neurons (Zeigler et al., 2002; Ponzio et al., 2006; Romanov et al., 2017). Third, neurons in the mammalian paraventricular hypothalamus heavily innervate the pre-locus coeruleus (region adjacent to the noradrenergic locus coeruleus), (Geerling et al., 2010), suggesting an anatomical correspondence between the preoptic-to-RoL1 projection we observe in zebrafish; both projections originate from homologous regions of the neurosecretory hypothalamus and project to a catecholamine-negative set of neurons just adjacent to the locus coeruleus. These similarities between larval zebrafish and rodents suggest that our findings are relevant to the mammalian brain and may be a conserved feature of vertebrate hypothalamus-brainstem interactions.

We observed that RoL1 neurons are an essential component of the fast avoidance response to environmental stressors. These neurons are known to drive locomotor behavior; optogenetic activation around RoL1 in zebrafish promotes swimming actions (Vladimirov et al., 2018; Marquart et al., 2019). Therefore rapid activation of these cells by hypothalamic inputs can provide the means of promoting avoidance or escape. Our anatomical analyses indicated that some RoL1-projecting peptidergic neurons co-express the vesicular glutamate transporter *vglut2a*, and that application of an AMPA receptor blocker around RoL1 neurons prevents the avoidance response to increases in salinity or acidity. However, further studies are required to determine whether these peptidergic cells use glutamate co-release to activate RoL1 neurons, and what role, if any, is played by the release of the peptide itself into the hindbrain. An alternative scenario is that different neuropeptides converge on a common excitatory intracellular signaling pathway in RoL1 neurons downstream of their specific metabotropic receptors, which could function in a degenerate manner (Marder, 2012; Nath et al., 2016). Given the prevalence of neuromodulator/amino-acid co-release throughout nervous systems (Schöne & Burdakov, 2012; Granger et al., 2017; Nusbaum et al., 2017), resolving these issues will be generally important for studies of neural circuit function across species and behaviors.

Our brain-wide and hypothalamic cell type-specific imaging data both point towards a similar principle: neural activity patterns are widely distributed across cell types and brain areas, even during simple behaviors. These types of widespread activity patterns have been observed in other studies that use large-scale cellular activity recordings across the nervous system (Wu et al., 1994; Ahrens et al., 2012; Kato et al., 2015; Lovett-Barron et al., 2017; Allen et al., 2019). While it is unlikely that every neuron active during a given behavior is required for its execution, it is not known whether such distributed activity is an incidental consequence of dense connectivity or reflects an important role in enforcing the reliability of common behaviors (Edelman & Gally, 2001). Here we found that multiple peptidergic neurons could drive a common avoidance behavior in response to various stressors, but that none played an essential role; this type of overlapping, potentially degenerate organization may be a common feature of hypothalamic circuits that control essential survival behaviors (Betley et al., 2013). More complex functions such as social recognition, comparison of need states, and decision making may require more intricate activity patterns among hypothalamic cell types (Lin et al., 2011; Burnett et al., 2016; Remedios et al., 2017). Under these conditions, functionally similar groups of neurons could be categorized differently according to their neuronal activity and behavioral contributions across different timescales and conditions, potentially revealing an organization distinct from cell type categorization through static anatomical and gene expression properties. These principles can be explored explicitly in future studies by recording from, and manipulating, multiple distinct molecularly-defined cell types on long timescales across behaviors of varying complexity.

Our results fit within a broader field of work that has established the hypothalamus can drive behaviors on a much faster timescale than previously appreciated, including fighting, mating, hunting, and escape (Lin et al., 2011; Lee et al., 2014; Füzesi et al., 2016; Li et al., 2018; Wang et al., 2019). Our work contributes to these efforts by demonstrating that neurosecretory cell types in the vertebrate hypothalamus are capable of generating avoidance behavior in response to stressors, through a convergent hypothalamus-brainstem-spinal pathway. We speculate that neurosecretory populations may be able to support their multiple roles by initiating slow specific adaptations through peptide secretion in the pituitary, and fast generalized avoidance through synaptic glutamate release in the hindbrain. These dual functional roles could be achieved through two populations of neurons (such as magnocellular and parvocellular neurons) or by diverging projections from the same neurons.

Neurosecretory cell types such as hypothalamic peptidergic neurons are hypothesized to be among the earliest neuron types to emerge during animal evolution (Hartenstein, 2006). Primitive secretory cells may have adopted multiple functional roles early in the evolution of nervous systems (Tessmar-Raible, et al., 2007), including the establishment of synaptic outputs to promote behavior on faster timescales than systemic hormone and peptide secretion allows. These multifunctional properties also appear to be present in the neurosecretory cells and circuits of the vertebrate hypothalamus: each hypothalamic “cell type” can play multiple roles across different timescales and behavioral circumstances, and multiple cell types can also serve common roles. This flexible, overlapping organization may ensure robust and reliable execution of the critical survival functions of the hypothalamus, an essential component of vertebrate life.

## Acknowledgements

We thank our colleagues for generously sharing transgenic zebrafish lines: Adam Douglass, Herwig Baier, Misha Ahrens, Suresh Jesuthasan, Josh Bonkowsky, and Jiu-lin Du. We thank Christine Constantinople, Felicity Gore, Tim Machado, Will Allen, Ravi Nath, and Claire Bedbrook for feedback on the manuscript, and the entire Deisseroth lab for advice and discussions. This research was supported by Helen Hay Whitney Foundation Postdoctoral Fellowships (MLB and ASA), Brain and Behavior Research Foundation NARSAD Young Investigator Awards (MLB and ASA), NIMH (MLB: K99MH11284002), NIDA (SG: R01DA035680 and R21DA038447), and grants from the NIH, NSF, and NOMIS Foundation (KD).

## Author contributions

MLB and KD designed experiments. MLB, RC, and SB performed experiments. MLB analyzed data. RC developed *in situ* hybridization protocols. ASA contributed custom software. MW and SG contributed transgenic zebrafish. MLB and KD wrote the paper with input from all authors. KD supervised all aspects of the work.

## METHODS

### Key Resources Table

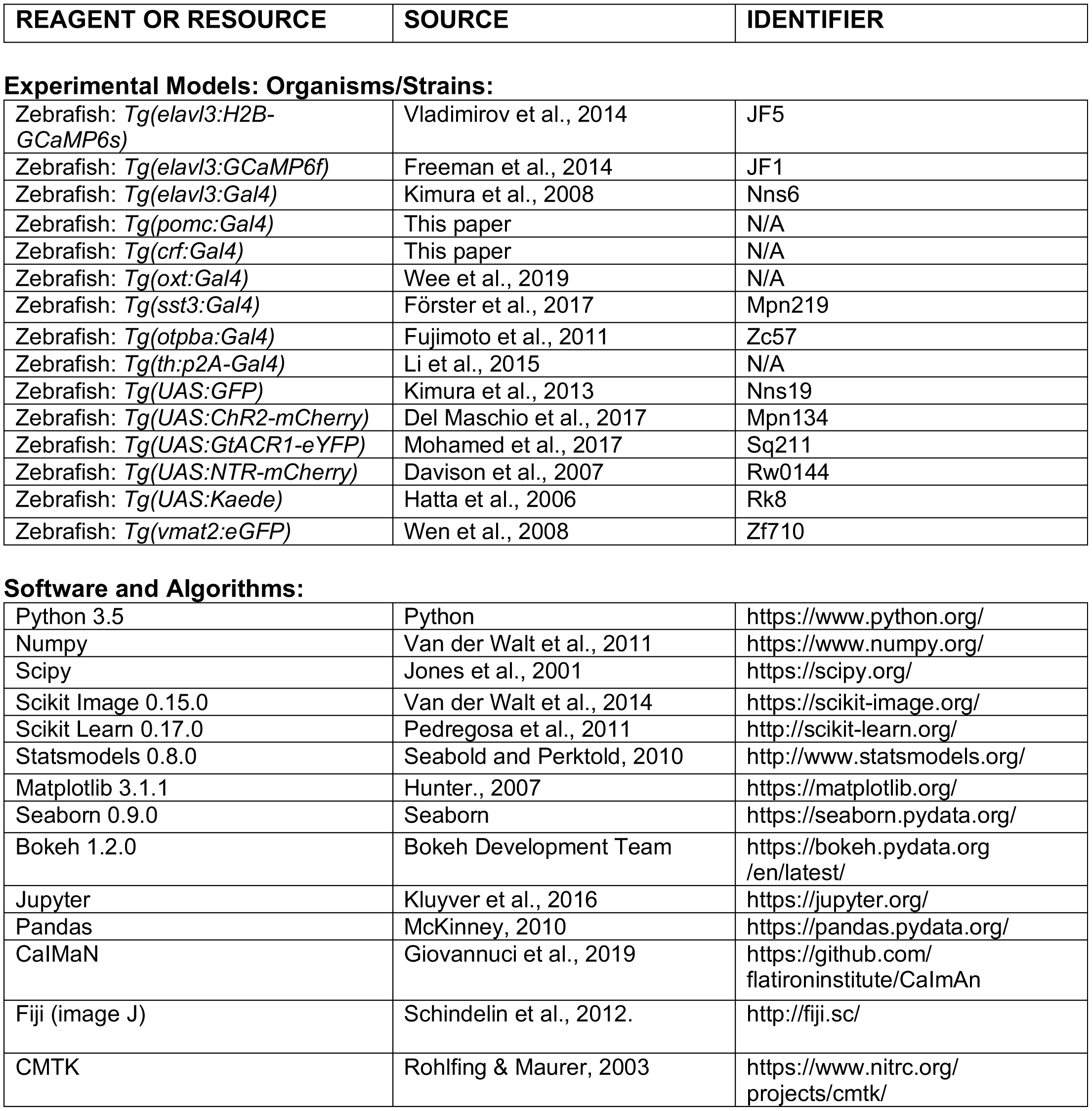

## EXPERIMENTAL MODEL AND SUBJECT DETAILS

All procedures were approved by the Stanford University Institutional Animal Care and Use Committee.

## METHOD DETAILS

### Zebrafish

We used larval zebrafish for all experiments in this study, tested at 6-8 days post fertilization (dpf). Animals were group-housed in a standard 14-10 h light-dark cycle, temperature-controlled room and raised according to Zebrafish International Resource Center (ZIRC) guidelines. All experiments were conducted during the light period (9am-6pm). At these stages of development, the sex of larval zebrafish is not yet defined. Larvae were fed with paramecia (Paramecia Vap Co.) twice daily from 5 dpf onward. GCaMP imaging was performed with homozygous fish, but otherwise fish were heterozygous for each transgene. We used the following published lines: *Tg(elavl3:H2B-GCaMP6s)* (Vladimirov et al., 2014), *Tg(elavl3:GCaMP6f)* (Freeman et al., 2014), *Tg(elavl3:Gal4)* (Kimura et al., 2008), *Tg(oxt:Gal4)* (Wee et al., 2019), *Tg(sst3:Gal4)* (Förster et al., 2017), *Tg(otpba:Gal4)* (Fujimoto et al., 2011), *Tg(th:p2A-Gal4)* (Li et al., 2015), *Tg(UAS:GFP)* (Kimura et al., 2013), *Tg(UAS:ChR2-mCherry)* (Dal Maschio et al., 2017), *Tg(UAS:GtACR1-eYFP)* (Mohamed et al., 2017), *Tg(UAS:NTR-mCherry)* (Davison et al., 2007), *Tg(UAS:Kaede)* (Hatta et al., 2006), and *Tg(vmat2:eGFP)* (Wen et al., 2008). For the *Tg(crf:Gal4)* line, upstream regulatory regions along with first intron of the *crhb* locus was used to drive the expression of Gal4. F1 larvae with hypothalamic expression overlapping endogenous *crf* were raised to establish the line. Detailed characterization of the line will be published elsewhere. The *Tg(pomc:Gal4)* line was generated using promoter elements described elsewhere (Liu et al., 2003).

### Behavioral Experiments

Zebrafish were mounted dorsal side up in a thin layer of 2.5% low-melting point agarose (Invitrogen) in the lid of a 3 mm petri dish (Fisher), using a sewing needle to position the fish under a stereomicroscope (Leica M80). After agarose solidified (10-15 minutes), agarose was removed from around the tail posterior to the pectoral fins and around the nose and mouth with a fine scalpel. Fish were then habituated next to the experimental apparatus for at least 20 minutes in E3 solution at room temperature (5 mM NaCl, 174 µM, KCl, 396 µM CaCl_2_, 673 µM MgSO_4_, 1 mM HEPES, with 1 M KOH added to pH = 7.2), before beginning behavioral testing. During behavioral testing, fish were exposed to a constant flow of E3 solution at 1.5 mL per minute through a 200 µm opening at the end of a pipette tip, positioned at 20-30 ° in front of the fish (pointed at the nose and mouth) with a micromanipulator (WPI, M3301L). Flow switched between E3 solution, increased salinity (50 mM NaCl added to E3), increased acidity (HCl (∼0.1 mM) or citric acid added to E3 to pH 4.8), or blank (E3), using solenoid pinch valves (Cole Palmer, P/N98302-02) controlled by a digital I/O device (National Instruments, USB-6525). Heat stimuli were delivered by a 980 nm DPSS Laser (Changchun New Industries Optoelectronics Technology Co, Ltd, MDL-III-980), through a 400 µm 0.48 NA fiber (Doric Lenses), threaded through the flow pipette tip and positioned 1 mm from the nose. This laser was controlled with the analog output of a multifunctional I/O device (National Instruments, USB-6003 or SCB-68A). Heat stimuli were 16 mW when measured from the fiber tip, and were measured as an increase of 7 ° C on a thermocouple (Warner Instrument Corp., TC-344B), in E3. Looming stimuli were projected at 60 Hz to a diffusive screen under the fish as three consecutive presentations of expanding black dots on a red background, using a Laser Pico Projector (MicroVision), surrounded by three Red Wratten filters (Kodak). Stimuli expanded with size increasing as constant approach velocity (0.05 mm to 3 mm over 3 s, using equations from Dunn et al., 2016). The tail of the fish was illuminated by IR lights from above and behind the fish, and tail movements were filmed at 60 frames per second from below with an AVT Manta G031 camera (Allied Vision) through a AF-S DX Micro Nikkor 85 mm f/3.5G ED VR macro lens (Nikon). Stimulus generation and behavioral recording were coordinated via custom Python software. This configuration was used for stand-alone behavioral experiments, and experiments under the two-photon microscope. Each fish was exposed to five trials. Each trial was 5 minutes in duration, with constant flow at 1.5 mL/min: 60 s baseline, 40 s salinity, 30 s baseline, 40 s acidity, 30 s baseline, 40 s heat, 30 s baseline, 9 s looming stimuli, 21 s baseline.

For ChR2 stimulation experiments, fish were embedded as described above, but without the agarose removed from around the nose. Neurons were stimulated with a 473 nm DPSS laser (OEM laser system Inc, BL-473-00100-CWM-SD-05-LED-0), through a 105 µm 0.22 NA fiber (Doric Lenses), positioned between the eyes at ∼80-90 ° with a micromanipulator (WPI, M3301L), making slight contact with the agarose overlying the fish. Light was delivered as 100 ms pulses at 5 Hz for 10 seconds (30-40 s inter-trial interval). For optogenetic inhibition experiments, the same 5-trial behavior was used as above, and constant blue light (8 mW measured from fiber tip) was delivered on trial # 2 and 4 (of 5) for the entire duration of each stimulus, directed between the eyes. Optogenetic localization was estimated with photoconversion of kaede from green to red, using 405 nm light (Thorlabs, M405FP1) directed through the optical stimulation fiber between the eyes of a *Tg(elavl3:Gal4; UAS:Kaede)* fish.

For drug injection experiments, *Tg(elavl3:H2B-GCaMP6s)* fish were embedded in agarose with the nose and tail free to move. Fish were transiently anesthetized by cooling and visualized with an upright microscope (Olympus BX51WI) equipped with DIC optics, and filter sets for visualizing Texas red and GCaMP. Glass pipettes were used to make a small incision at the fissure between the optic tectum and lateral cerebellum, approached from the side between the ear and eye, on both sides of the fish. A glass pipette with a ∼5-10 µm diameter was filled with either NBQX (Tocris, 25 µM in PBS, 1:2000 DMSO, Texas red dye for visualization) or vehicle (PBS, 1:2000 DMSO, Texas red dye for visualization) and inserted into the brain, starting in the fissure between the tectum and lateral cerebellum. We used a fresh pipette for each fish, to minimize clogging. Under visual guidance the pipette tip was placed around the RoL1 cell bodies, lateral to the raphe nucleus (visualized with H2B-GCaMP label), and drug was pressure ejected. Injection extent was monitored by observing Texas red fluorescence, and the pipette was removed once the RoL1 region was covered in fluoresence. The fish was then rotated and injected on the other side. Fish were allowed to recover for 5-10 minutes before behavior initiated. All fish were confirmed to be capable of movement, by checking for tap-evoked startle.

### Two-photon Ca^2+^ imaging

Two-photon Ca^2+^imaging was performed with an Olympus FVMPE multiphoton microscope (Olympus Corporation), with a resonant scanner in bidirectional scanning mode. For brain-wide live imaging at 920 nm, we used a 16x objective (0.8 NA; Nikon) at 1.1x zoom in 14 z-planes, cropped to 512×305 pixels, separated by 15 µm, at 2 volumes/second (2800 volumes). After completion of behavior and functional brain imaging, a structural stack was obtained at 820 nm, with 1 µm spacing and 16x frame averaging, starting 15 µm above the first z-plane, ending 15 µm below the last z-plane, and repeated twice. For hypothalamus imaging at 920 nm, we used a 25x objective (1.05 NA; Olympus) at 2x zoom in 5 z-planes, cropped to 5123×05 pixels, separated by 15 µm, at 4 volumes/second (6500 volumes). After completion of behavior and functional brain imaging, a structural stack was obtained at 820 nm, with 1 µm spacing and 16x frame averaging, starting 15 µm above the first z-plane, ending 15 µm below the last z-plane, and repeated twice. For SPN imaging, we used a 25x objective (1.05 NA; Olympus) at 1.2x zoom in 6 z-planes, cropped to 512×305 pixels, separated by 20 µm, at 4 volumes/second (5500 volumes).

### Fluorescent *in situ* hybridization

To eliminate the need for probe optimization and suppress background signal, we designed hybridization probes according to the split initiator approach of third-generation *in situ* hybridization chain reaction HCR v3.0 (Choi et al., 2018).

Even and odd 22-25 nt DNA antisense oligo pairs carrying split B1, B3 or B5 initiation sequences were tiled across the length of the mRNA transcript, synthesized by IDT (Integrated DNA Technologies, Inc), and used without further purification. Dye-conjugated hairpins (B1-647, B3-546, and B5-405) were purchased from Molecular Technologies (Pasadena, CA).

Zebrafish were fixed overnight in 4% PFA in 1X PBST at 4 °C. After washing (3 times in 1X PBST; 5 m each), larvae were permeabilized for 10 m in 100% (v/v) methanol at -20 °C and then rehydrated (50% (v/v) methanol, 25% (v/v) methanol, then in 2X SSCT; 5 m each). Hybridization with split probes were performed overnight in 2X SSCT, 10% (v/v) dextran sulfate, 10% (v/v) formamide at 4 nM probe concentration. The next day, larvae were washed (3 times in 2x SSCT, 30% (v/v) formamide at 37 °C then 2 times in 2X SSCT at room temperature; 20 m each) then incubated in amplification buffer (5X SSCT, 10% (v/v) dextran sulfate). During this time, dye-conjugated hairpins were heated to 95 °C for 1 m then snap-cooled on ice. Hairpin amplification was performed by incubating individual zebrafish in 50 µL of amplification buffer with B1, B3, and B5 probes at concentrations of 240 nM overnight in the dark. Samples were washed 3 times with 5X SSCT for 20 m each, then samples were mounted in 2-3% low-melting point agarose, covered in 2x SSCT or PBS, and immediately imaged under the two-photon microscope. All four channels were imaged simultaneously, in unidirectional resonant scanning mode (16x line average), at 820 nm.

To perform multi-round HCR v3.0, larvae were excised and kept in agarose blocks and digested overnight in DNase I (0.2 units/µL) at room temperature. Following 3 rounds of washing with 2X SSCT for 1 h each, hybridization and amplification were performed with the same protocol as the first step. We performed multi-round HCR-FISH with the following combination of probe sets: (Round AVP-B1/CRH-B3/OXT-B5; (Round 2) NPY-B1/VIP-B3/SST-B5; (Round 3) HCRT-B1/NPVF-B3/TRH-B5. The list of probes (*avp, oxt, crf, sst, npy, vip, hcrt, npvf, trh, vglut2a*) is found in Supplementary Table 1.

For counting neurons that are dual *vglut2a*^+^ and peptide^+^, four images of the preoptic hypothalamus were collected from two animals in each group, from top-down two-photon scans at 820 nm. Neurons were counted from 100 µm x 100 µm areas around the third ventricle. Neuron counts for each sample are (# *vglut2a*^+^peptide^+^ / # peptide^+^): *Tg(otopba:Gal4; UAS:GFP)*): 25/27 cells, 12/13 cells, 5/7 cells, 13/13 cells (total = 55/60 cells); *oxt*: 5/8 cells, 20/20 cells, 16/17 cells, 10/10 cells (total = 51/55 cells); *crf*: 10/11 cells, 18/19 cells, 7/7 cells, 9/9 cells (total = 44/46 cells); *sst*: 0/4 cells, 1/11 cells, 2/12 cells, 1/7 cells (total = 4/34 cells).

### Volume registration

All volume registrations were performed in bright *Tg(elavl3:H2B-GCaMP6s)* fish. Image volumes were saved as .nrrd files in a mm scale. For brain-wide imaging experiments, all anatomical brain volumes were registered to a common “bridge brain” volume from an exemplar fish. Using the Z-Brain Atlas (Randlett et al., 2015), the “*Tg(elavl3:H2B-RFP)”* volume and each Z-brain mask (294 in total) were registered to the bridge brain. By moving all neuron coordinates to the same common volume, the identity of neurons in each region could be established with Z-brain mask overlap. For MultiMAP experiments, volumes of endogenous GCaMP fluorescence from each round of fluorescent *in situ* hybridization were registered to the live GCaMP anatomical volume in the same fish to match the same cells in both volumes.

Registration was achieved using the Computational Morphometry Toolkit (CMTK; Rohlfing and Maurer, 2003), using initial affine, affine, and b-splines registration steps (Lovett-Barron et al., 2017). Volumes were registered using CMTK installed on Amazon Web Service’s cloud computing environment (c3.8xlarge instances). Once the final transformation was determined, the transformation coordinates were applied to the fixed GCaMP volume and each of the associated fluorescent *in situ* hybridization volumes. From each of these volumes now aligned to the live GCaMP volume, z-planes were extracted that correspond to the z-planes with activity recorded (every 15 µm, from 15 µm below the dorsal extent and 15 µm above the ventral extent). Each z-plane in each fish was manually inspected to ensure cellular-resolution registration, before selection of labeled neurons in these planes.

Neurons were selected using custom software in Python, using Bokeh for interactive image processing in a Jupyter notebook. The total number of peptidergic neurons (ordered as [*avp, oxt, crh-medial, crh-lateral, npy-dorsal, npy-ventral, vip-anterior, vip-posterior, sst*]) identified for each of the 7 fish were: [4, 6, 8, 4, 7, 17, 7, 12, 5], [3, 10, 2, 7, 16, 5, 3, 8, 6], [5, 6, 3, 4, 5, 4, 5, 10, 12], [8, 7, 5, 7, 13, 4, 3, 12, 11], [4, 5, 7, 9, 10, 6, 3, 7, 11], [4, 16, 7, 10, 14, 9, 6, 11, 10], and [5, 3, 5, 3,10, 3, 1, 11, 8]. Many *avp*^+^ neurons were also *crh*^+^, as has been previously reported (Herget et al., 2015).

### Neuron Ablation

#### NTR ablation

Fish were sorted for NTR-mCherry expression using an upright fluorescence dissecting microscope (Leica, M165 FC). mCherry^+^ fish and an equal number of pigment-matched mCherry-clutch mate controls were collected, and placed into dishes with E3 containing 10 mM metronidazole (MTZ, MP Biomedicals, 02155710) at 4 dpf, and provided with a drop of paramecia. Dishes were covered in tin foil, and removed 36 hours later, at 5.5 dpf. Fish were then inspected to ensure decreased or absent mCherry fluorescence before continuation of experiment. Fish were transferred to standard E3, provided with paramecia, and allowed to recover until testing at 7 or 8 dpf (interleaving control and NTR^+^ fish over the experimental session). Subsets of fish were preserved in PFA for subsequent anatomical imaging. Some *Tg(otpba:Gal4; UAS:NTR-mCherry)* fish showed a distended swim bladder after ablation, as has been previously noted (Lambert et al., 2012). However, these fish were still capable of spontaneous locomotion and large-angle turns during agarose fixation (Figure S5d), unlike the locomotor phenotypes observed upon ablation of these neurons at earlier stages – a consequence of decreased descending dopaminergic signaling during development (Lambert et al., 2012).

#### Two-photon ablation

For spinal projection neuron labeling, Texas Red Dextran (10000 mW, lysine fixable; Invitrogen, D1863) was injected into the spinal cord of 5 dpf *Tg(elavl3:H2B-GCaMP6s)* fish anesthetized with 0.1% MS-222 (Sigma) and fully embedded in agarose. Fish were cut out of agarose and placed in normal fish system water to recover for 24 hours. At 6 dpf, successfully labeled fish were mounted in agarose, and placed under the 2-photon microscope. Relevant Texas Red^+^ neurons were identified based on morphological properties, and were ablated using galvo scanning to focus 780 nm to a small excitation volume around nucleus. Power was manually increased while monitoring GCaMP and Texas Red fluorescence. Ablation was halted once GCaMP fluorescence sharply increased and Texas Red fluorescence decreased (typically 2-8 seconds). After each cell was ablated, an anatomical image was taken to monitor the localization and extent of the ablation. If the cell remained, this procedure was repeated until the cell was ablated. If the damage exceeded the extent of the single cell targeted, the fish was not used for subsequent experiments. Sham ablation fish were treated to the same procedure, but imaged at 780 nm without sufficient power to ablate neurons. Successfully ablated and sham ablation fish were cut out of agarose, and returned to a petri dish with E3 and provided with paramecia. Fish were tested 24 hours later at 7 dpf.

## QUANTIFICATION AND STATISTICAL ANALYSIS

All analyses and visualizations were performed with custom code written in Python, using NumPy, Scipy, Matplotlib, Jupyter, Seaborn, Statsmodels, Pandas, Scikit-image, and Scikit-learn libraries (Jones et al., 2001; Hunter, 2007; McKinney, 2010; Kluyver et al., 2016; Seabold & Perktold, 2010; Pedregosa et al., 2011; van der Walt et al., 2011; van der Walt et al., 2014).

### Behavioral analysis

The pixels containing the fish tail were determined in each frame of the tail monitoring videos using adaptive thresholding and blob detection algorithms in Scikit-image (Andalman et al., 2019, Lovett-Barron et al., 2017). Tail movements were identified by analyzing the number of tail-containing pixels that did not overlap between adjacent frames. The mean and standard deviation of this value when the tail was motionless was estimated as the median of these statistics within all 300 ms time bins. Tail movements were identified as frames when this value remained 4 standard deviations above baseline for at least 40 ms. Tail movements that were separated by less than 50 ms were merged. To classify movement types, the orientation of the tail was computed for every frame as the angle between neutral tail position and the major-axis of the ellipse fit to second-moment of the pixels containing the tail. Movements were then classified as turns or escapes if the maximum angle of deflection exceeded 30 degrees and the maximum velocity of deflection exceeded 1.5 degrees/ms. These movements were also classified in terms of their peak tail angle (95th percentile of all measured angles within movement). Further analyses were conducted on a binary array that spanned the entire behavioral session, with the times of turns/escape onsets or forward swimming onsets noted as ones. For display purposes in Figure 1b and Figure S1b, arrays were converted to 1 s bins of movement rate, spanning 10 s before stimulus onset to 10 s after stimulus offset (60 s total).

Stimulus-driven movement was defined as any turn/escape movement with an onset time within 40 s of stressor/stimulus (heat, salinity, acidity, or loom) onset. For each fish, the fraction of trials with movement was determined for each stimulus (5 trials of each), as well as the peak tail angle of any successful movements to each stimulus. For drug injection experiments we used the first 4 trials, because we observed drug washout by the 5^th^ trial. The same metrics were also determined for light-induced movement in ChR2 experiments, but the time frame analyzed was 2 s after light onset until 2 s after light offset (10 s total).

### Two-photon Ca^2+^ imaging data processing

Ca^2+^ imaging data were processed using the CaIMaN pipeline (Giovannuci et al., 2019). We performed piecewise motion correction in 128×128 pixel patches with a 48×48 pixel overlap. Source extraction was performed with an expected neuron size of 4×4 pixels (zoomed imaging) or 3×3 pixels (whole-brain imaging), using the ‘greedy_roi’ method, using patches of 50×50 pixels (initialized as 10 components per patch in zoomed imaging, and 20 components per patch in whole-brain imaging, merge threshold = 85%). The minimum signal-to-noise for accepting a component was 2.0, with a 90% pixel correlation. Deconvolved fluorescence traces (GCaMP decay = 3.0 s for nuclear-localized indicator, 1.0 s for cytosolic indicator) were then z-scored for further analysis. All behavioral and neural time series were aligned based on synchronized sampling of camera and microscope frame times, and resampled to a common 10 Hz sampling rate.

### Linear models of single neuron activity

Regressors were composed of boxcar functions matching behavioral features, and smoothed by an exponentially-weighted moving average with a decay of 3 s (approximate decay of fluorescence from H2B-GCaMP6s). The regressors were: array of turning/escape movement onsets, salinity stimulus (40 s), acidity stimulus (40 s), heat stimulus (40 s), and looming dot (9 s). In addition, the unsmoothed tail angle was used as a regressor. Five versions of each regressor were produced – the native timing, as well as shifted forward and backwards in time by 1 and 2 seconds. These 30 regressors (6 behavioral features x 5 time permutations) were used in the linear model.

Each neuron’s fluorescence time series was fit with a linear model of the regressors, using Elastic Net Regression in Scikit-learn (L1 ratio=0.1, 5 iterations, 5 alpha values, and 10x cross-validation), trained on 90% of the data and tested on 10% held out data. Note that the L1 ratio (scaling between L1 and L2 penalties, with 1.0 corresponding to Lasso regression) corresponds to the *alpha* variable in the glmnet package in R, whereas the alpha value here (a constant to multiply penalties) corresponds to the lambda variable in glmnet. After determining the fit of the full model (R^2^), each set of regressors (all time permutations) were shuffled, and this partial model was fit again. Unique model contributions (UMCs) for each task component were defined as the decrease in R^2^ from the full model to each partial model (Musall et al., 2018), and plotted as the percentage of the full model (Engelhard et al., 2019). Neurons with improvements in fits after shuffling had very low R^2^ from the full model, and were excluded from further analysis. Neurons were clustered by this UMC array, using spectral clustering in Scikit-learn (30 clusters, 50 nearest neighbors, *k*-means label assignment).

### Stimulus classification from population activity

In each fish, populations of neurons in each brain region were defined by molecular identity (MultiMAP experiments), or overlap with a Z-brain region, excluding sensory ganglia, overview regions (“Diencephalon –”, “Mesencephalon –”, “Rhombencephalon –”, and “Telencephalon –”), and regions with fewer than 20 neurons. A low-dimensional representation of each of these populations were obtained by principal component analysis (10 components). For each trial, z-scored temporal components were binned at 2 s, and each time bin of 2 s (10 components) was used to predict trial category (heat, salinity, acidity, loom), using a one-versus-rest classifier (Linear Support Vector Classification) trained on 80% of the trials and tested on 20% held out trials. Brain regions were ranked as the difference in the mean classification accuracy between the time of stimulus onset (the 40 s from stimulus onset to stimulus offset) minus the classification accuracy 10 s before stimulus onset. This same analysis was performed for behavior classification from peptidergic neuron populations, except that the traces of individual neurons were used instead of temporal principal components.

### Statistics

Statistics were obtained using parametric tests if the samples were normally distributed. The distribution of each sample was tested with the Shapiro-Wilk test, and if any distribution tested showed p<0.05, non-parametric tests were used for all comparisons in the same dataset. The exact test type is reported when used in the main text and/or figure legends. These tests were performed using Scipy in Python. P-values were corrected for multiple comparisons with the Benjamini/Hochberg false discovery rate correction, using the Statsmodels package in Python.

---

**Supplementary Figure 1.**
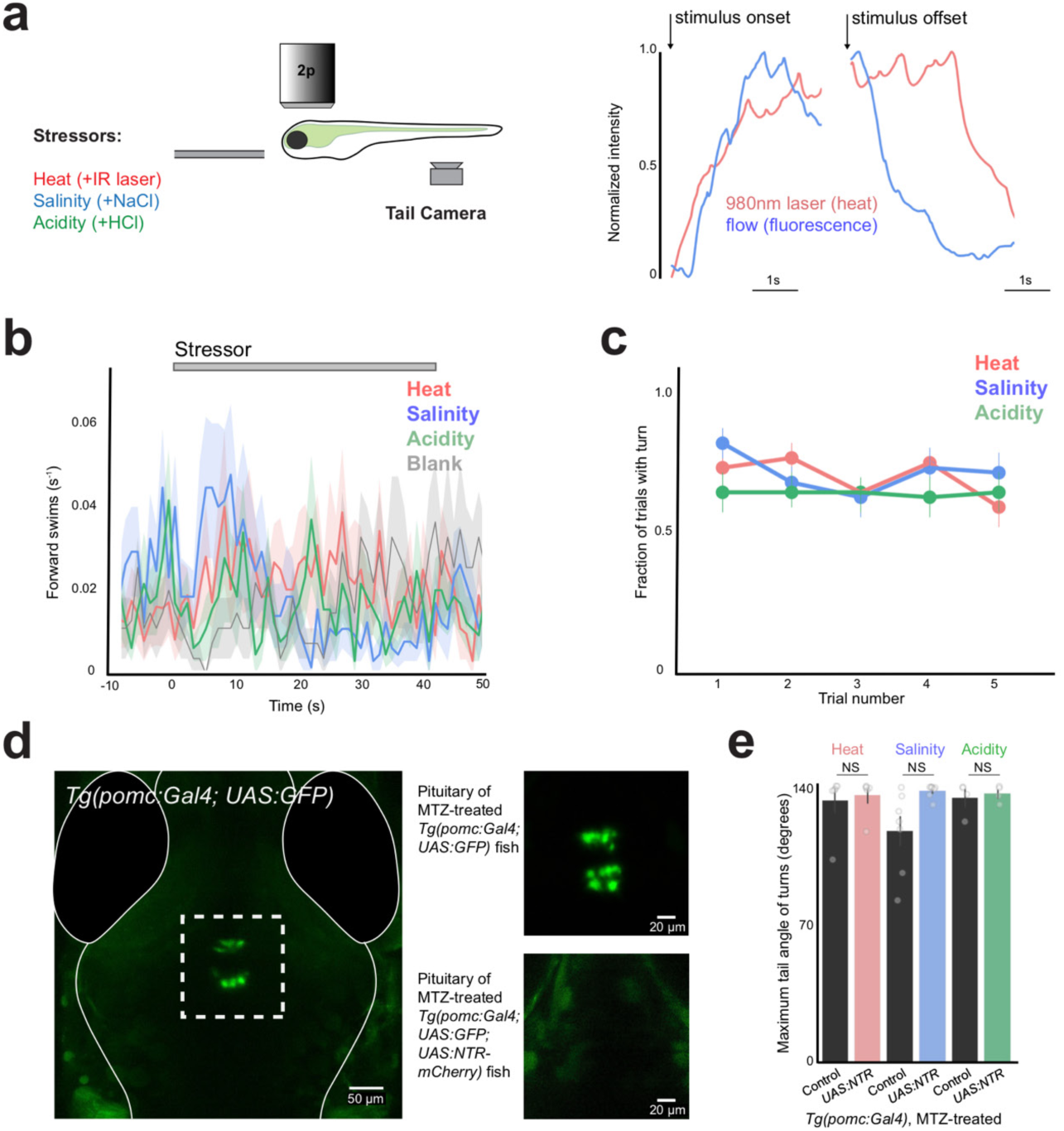
Rapid environmental changes do not influence forward swimming, responses do not habituate over trials, and ablation of *pomc*^+^ cells in the pituitary does not influence the magnitude of evoked turns during behavior. Related to Figure 1. **a)** Schematic of behavior (left), and measurement of stressor onset/offset kinetics using a thermocouple (heat) or two-photon imaging of fluorescence in flow solution (flow -salinity, acidity) (right). **b)** Stressor onset does not influence forward swimming (Mean ± SEM, 1 s bins, N = 57 fish (5 trials each), same fish as shown in Figure 1b). **c)** Fraction of trials with turning movements during stressor presentation, across all 5 trials (Mean ± SEM, N = 57 fish). Comparison of first two trials vs last two trials. All p> 0.1, Wilcoxon signed-rank test. **d)** Image of *Tg(pomc:Gal4;UAS:GFP)* fish, showing expression in pituitary corticotroph cells, with higher magnification image of control and ablated fish at right (co-expressing UAS:NTR-mCherry). **e)** Maximum tail angle of responsive movements - summary data from control and ablated fish. N = 9 fish per group, mean ± SEM, individual fish are points. Mann-Whitney U tests, correction for multiple comparisons. NS = “not significant”, all p > 0.4. Same fish as Figure 1d.

**Supplementary Figure 2.**
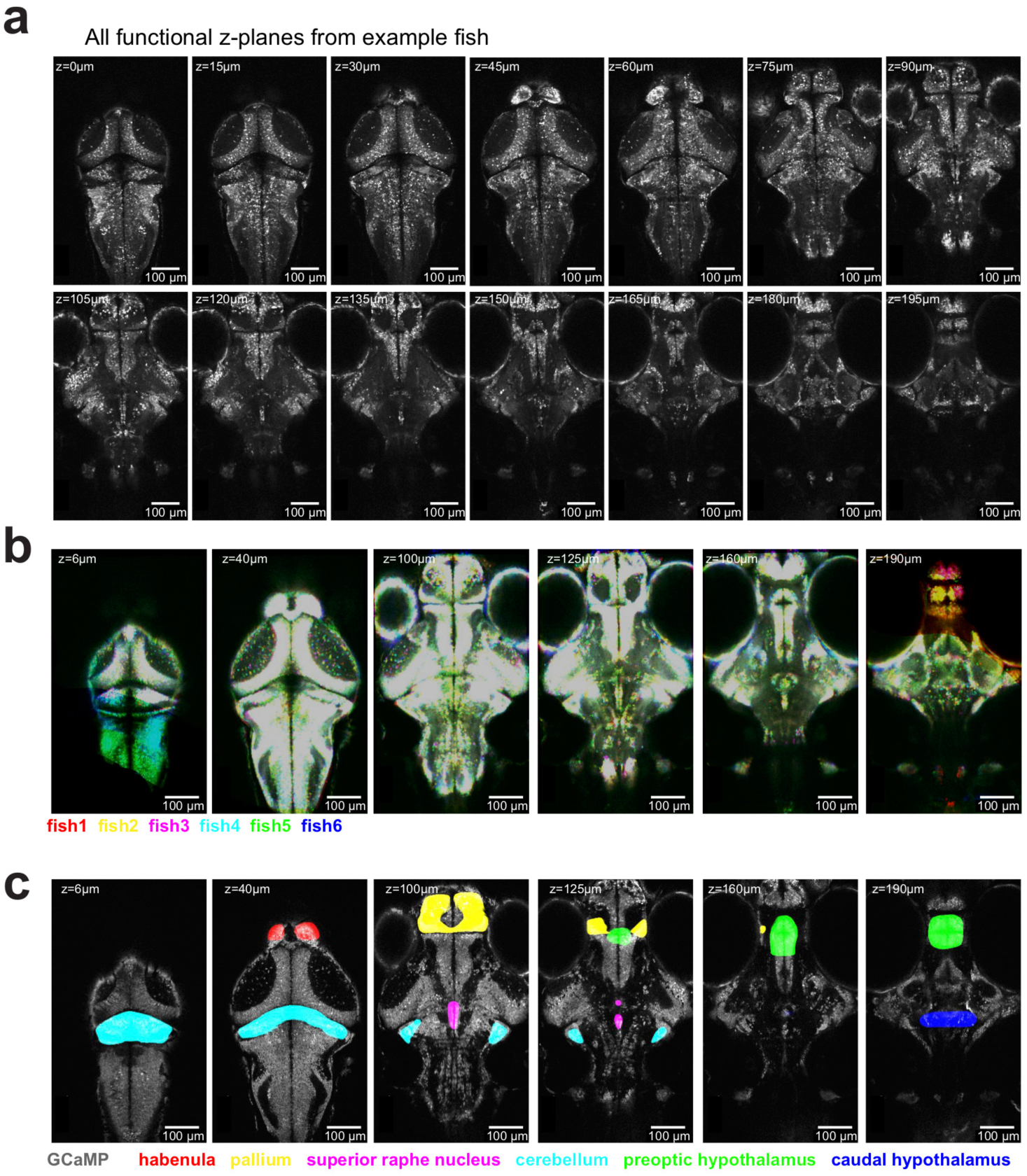
Examples showing coverage, registration, and anatomical segregation of brain-wide imaging data. Related to Figure 2. **a)** All z-planes functionally imaged in a single *Tg(elavl3:H2B-GCaMP6s)* fish - 14 z-planes, 15 µm separation, 2 volumes/second. **b)** Example z-planes from six fish, after registration to a common volume. Fish were registered to a common “bridge” brain; the standard *Tg(elavl3:H2B-RFP)* brain from the Z-Brain atlas (along with atlas masks) were registered to this bridge brain, to allow for anatomical region analysis. **c)** Example z-planes from a single fish, with a subset of overlaid anatomical regions from the registered Z-Brain atlas.

**Supplementary Figure 3.**
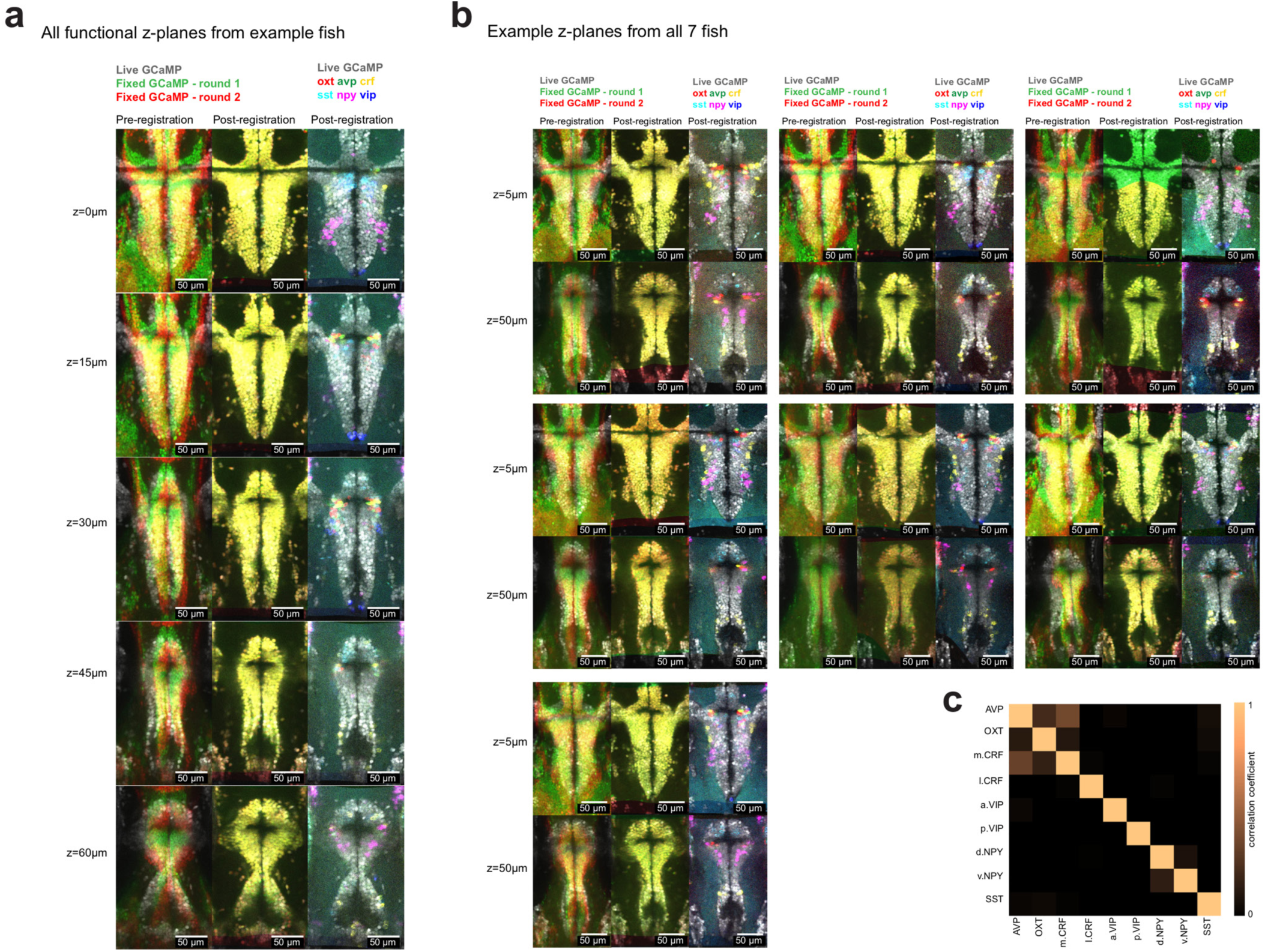
Cellular-resolution registration of live functional imaging to two rounds of multi-color fluorescent *in situ* hybridization. Related to Figures 3 and 4. **a)** All z-planes functionally imaged in a single *Tg(elavl3:H2B-GCaMP6s)* fish - 5 z-planes, 15 µm separation, 4 volumes/second. Showing live GCaMP z-plane (grey), and fixed GCaMP z-planes from round 1 and round 2 of *in situ* hybridization (green and red, respectively), before (left) and after (middle) volume registration. Overlap of live GCaMP and the six registered *in situ* hybridization labels from two rounds of multicolor fluorescent *in situ* hybridization (right). **b)** Example z-planes from all fish used in Figures 3 and 4. **c)** Overlap of molecular labels in cell types identified. Note the co-expression of *avp* with the medial *crf* population.

**Supplementary Figure 4.**
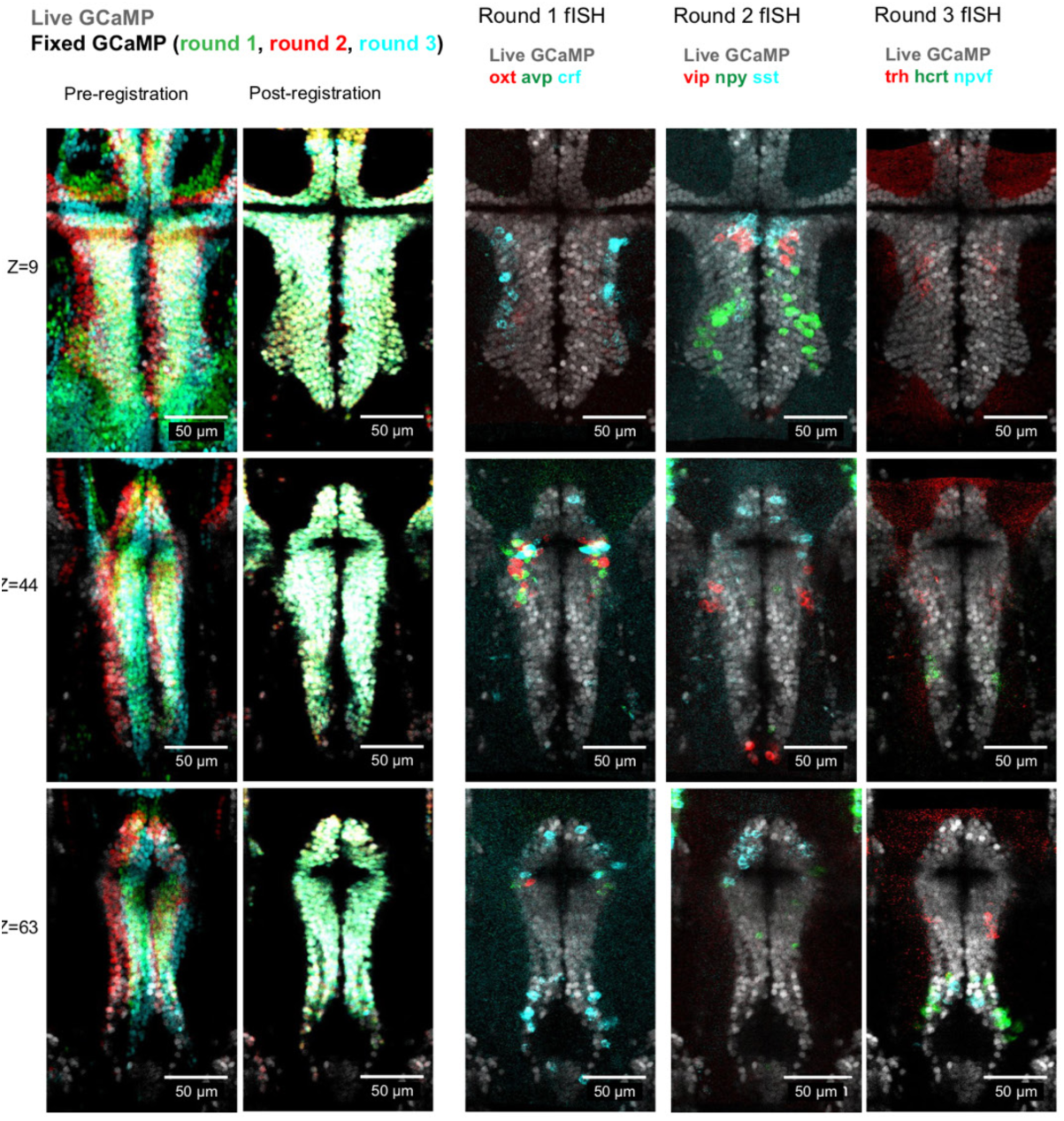
Example of cellular-resolution registration of live functional imaging to three rounds of multi-color fluorescent *in situ* hybridization - imaging nine peptidergic cell types at once. Related to Figures 3 and 4. Example z-planes functionally imaged in a single *Tg(elavl3:H2B-GCaMP6s)* fish, showing live GCaMP z-plane (grey), and fixed GCaMP z-planes from round 1, 2, and 3 of fluorescent *in situ* hybridization (green, red, and cyan, respectively), before and after volume registration. Overlap of live GCaMP and the nine registered *in situ* hybridization labels from three rounds of triple fluorescent *in situ* hybridization (right).

**Supplementary Figure 5.**
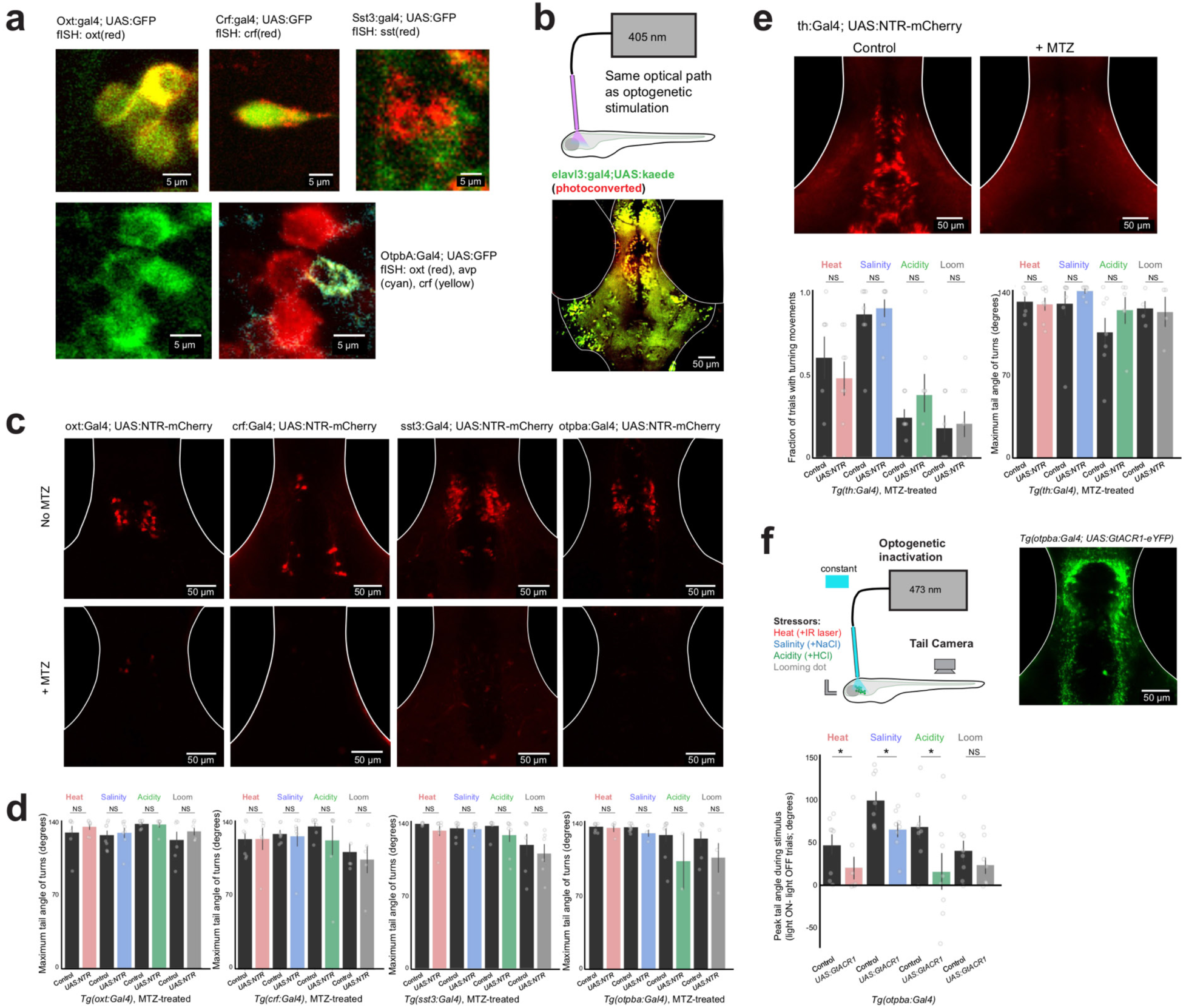
Characterization of Gal4 lines, optical excitation, and controls related to cellular ablation. Related to Figure 5. **a)** Overlap of GFP expression in the preoptic area of each Gal4 line, crossed UAS:GFP, with *in situ* hybridization labels, and overlap of GFP expression in *Tg(otpba:Gal4;UAS:GFP)* line with fluorescent *in situ* hybridization for *oxt, crf*, and *avp*. *Crf* and *avp* are co-localized in the same cells. **b)** Localization of optical stimulation for optogenetics experiments. 405nm light was delivered through the same optic fiber used for ChR2 or GtACR1 stimulation, and converts kaede from green to red. The increased green-to-red change in the preoptic area demonstrates limited spatial extent of optogenetic stimulation. **c)** Expression of UAS:NTR-mCherry in the preoptic hypothalamus of each Gal4 line, and its ablation upon treatment with MTZ (related to Figure 5e,f). **d)** Maximum tail angle of responsive movements - summary data from control and ablated fish. Same fish as Figure 5f. Mean ± SEM, individual fish are points. Mann-Whitney U tests, corrected for multiple comparisons within genotype. NS = “not significant”, all p > 0.1. **e)** Ablation of neurons in *Tg(th:Gal4)* line, including the dopaminergic posterior tuberculum neurons also labeled in the *Tg(otpba:Gal4)* line, does not influence behavior in this assay. N = 8 fish per group, mean ± SEM, individual fish are points. Mann-Whitney U test, corrected for multiple comparisons. NS = “not significant”, all p > 0.4. **f)** Inactivation of preoptic neurons in *Tg(otpba:Gal4; UAS:GtACR1-eYFP)* fish. Schematic of experiment (top), image of GtACR1-eYFP expression (middle), and behavioral results (bottom). N = 8 fish per group, mean ± SEM, individual fish are points. Mann-Whitney U tests, corrected for multiple comparisons. NS = “not significant”, all p > 0.1. *p < 0.05. While light ON trials increases total movement in control and opsin fish, GtACR1-expressing fish move less under these conditions.

**Supplementary Figure 6.**
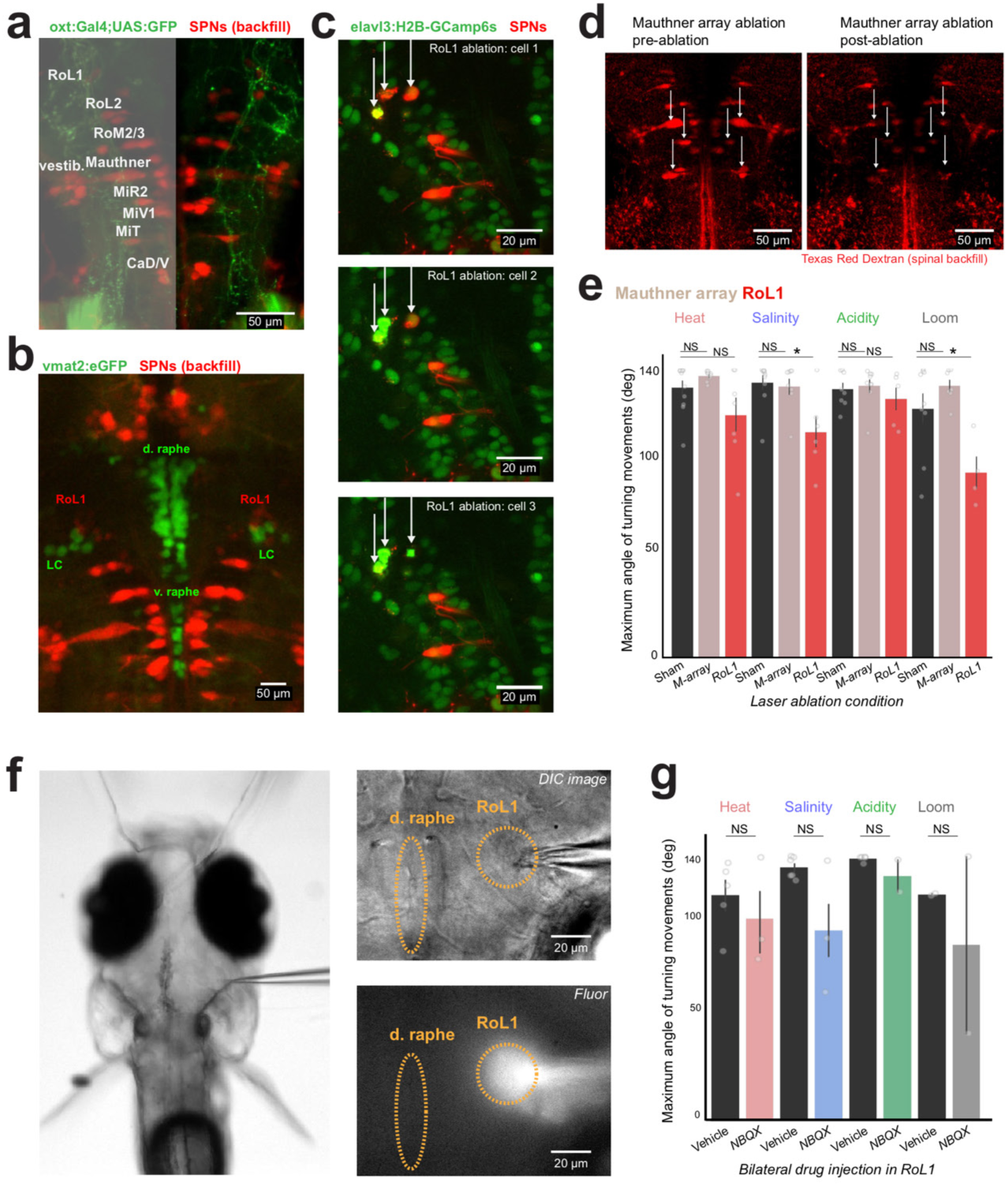
Details of RoL1 location, two-photon single-cell ablation of spinal projection neurons, and local injection of NBQX. Related to Figure 6. **a)** Location of spinal projection neurons (SPNs), with cell names, and axonal projections in *Tg(oxt:Gal4; UAS:GFP)* fish. **b)** Location of SPNs relative to monoaminergic cells, labeled with spinal backfill and *Tg(vmat2:eGFP)* line, respectively. RoL1 neurons reside slightly anterior and medial to the noradrenergic neurons of the locus coeruleus. **c)** Example images showing sequential two-photon single-cell ablation of three RoL1 neurons in *Tg(elavl3:H2B- GCaMP6s)* fish backfilled with Texas red dextran. **d)** Images of an example fish, backfilled with Texas red dextran, before and after bilateral ablation of the Mauthner neurons and segmental homologues (“M-array” ablation). **e)** Maximum tail angle of responsive movements - summary data from control and ablated fish. Mean ± SEM, individual fish are points. N = (11,11,10) – (sham, M-array, RoL1), left to right. Mann-Whitney U tests, corrected for multiple comparisons. Same fish as Figure 6e. NS = “not significant”, all p > 0.2. *p < 0.05. **f)** DIC image of embedded fish (agar around nose removed), with location of injection pipette entering from the fissure between the optic tectum and lateral cerebellum (left). Close-up image of injection area in DIC and fluorescence imaging to visualize drug/dye flow (right). **g)** Maximum tail angle of responsive movements - summary data from in RoL1-injected fish (NBQX or vehicle). Mean ± SEM, individual fish are points. N = (6,6) – (vehicle, NBQX). Mann-Whitney U tests, corrected for multiple comparisons. Same fish as Figure 6i. NS = “not significant”, all p > 0.1.

**Supplementary Table 1.**
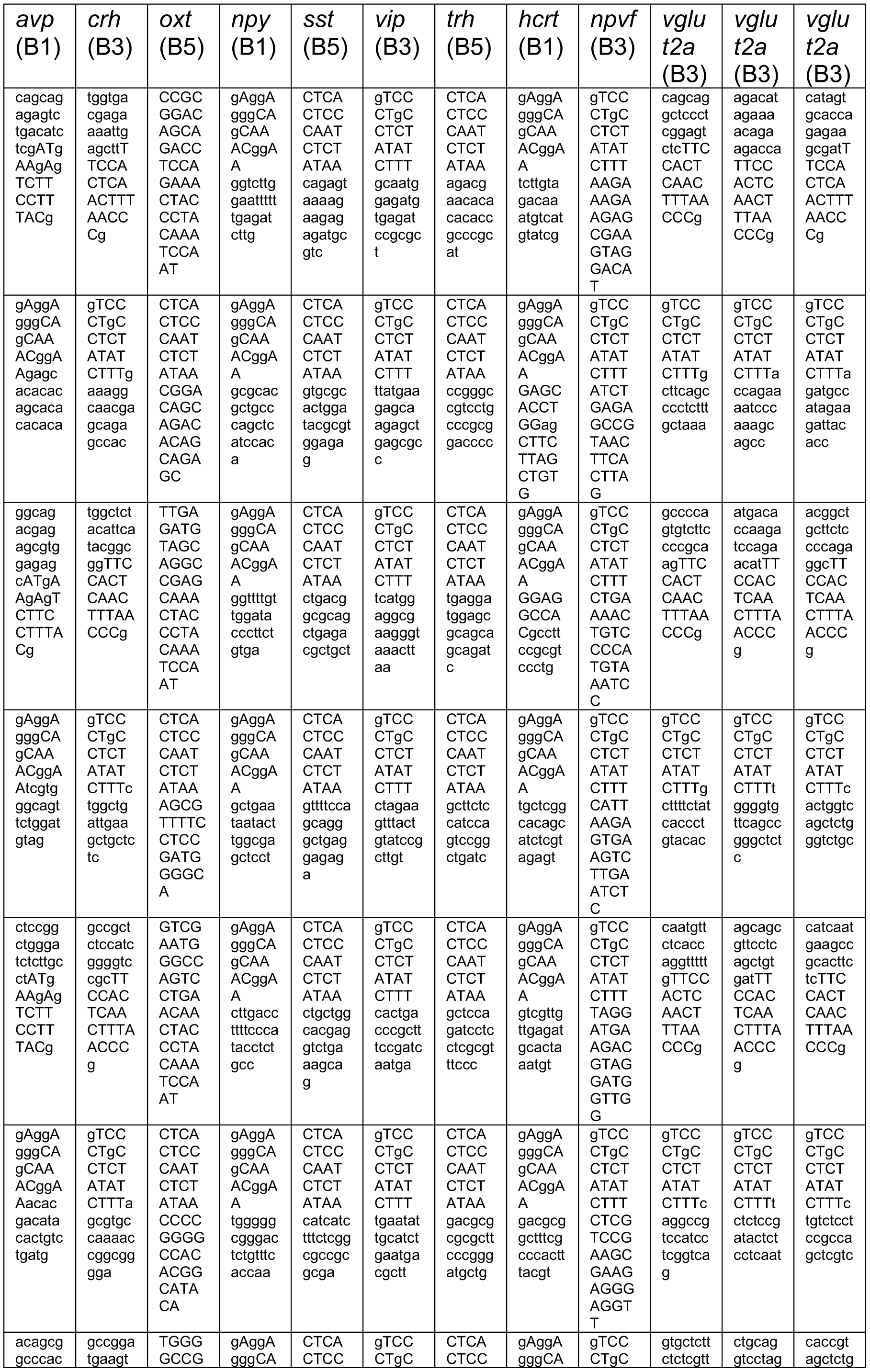
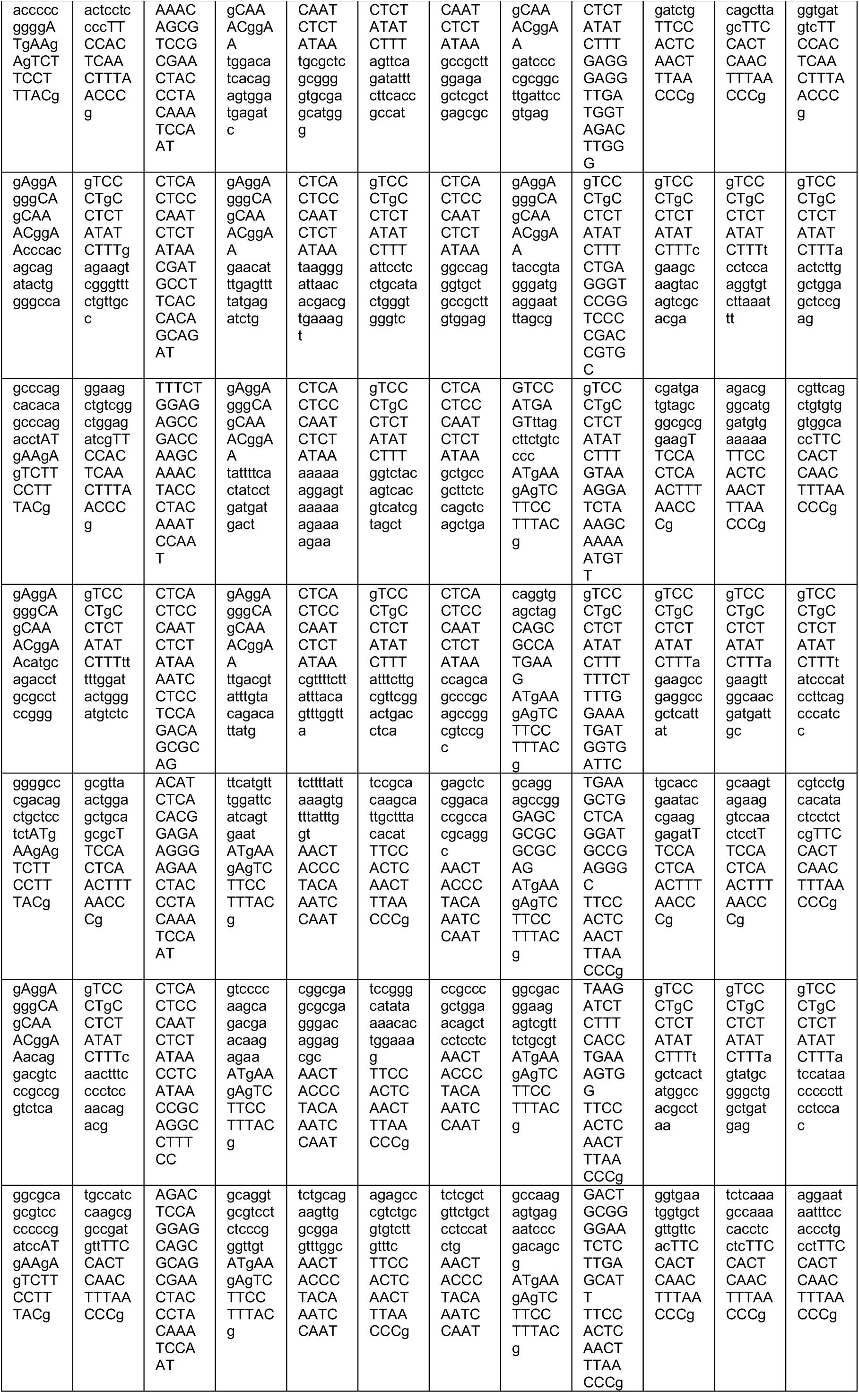
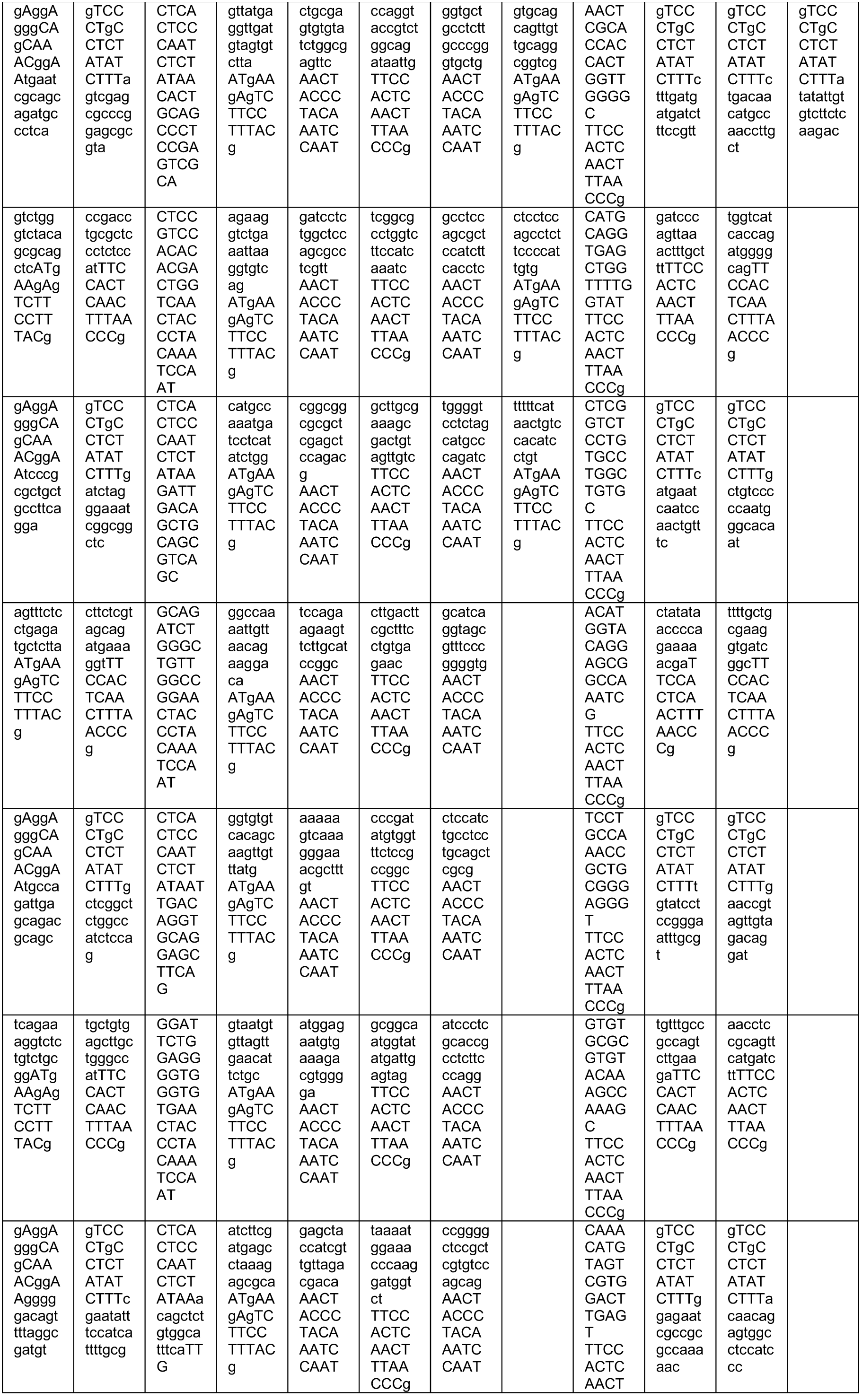
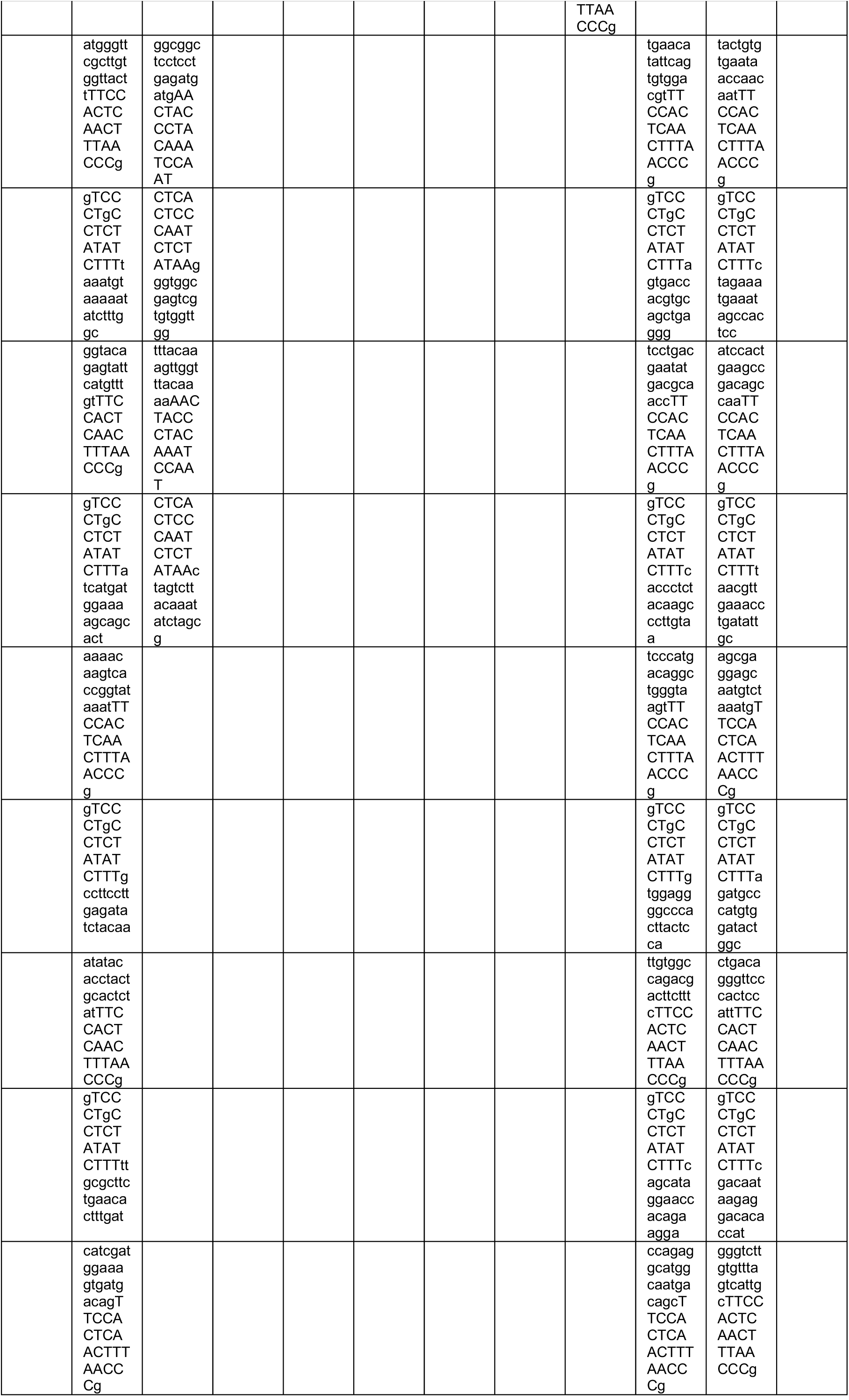
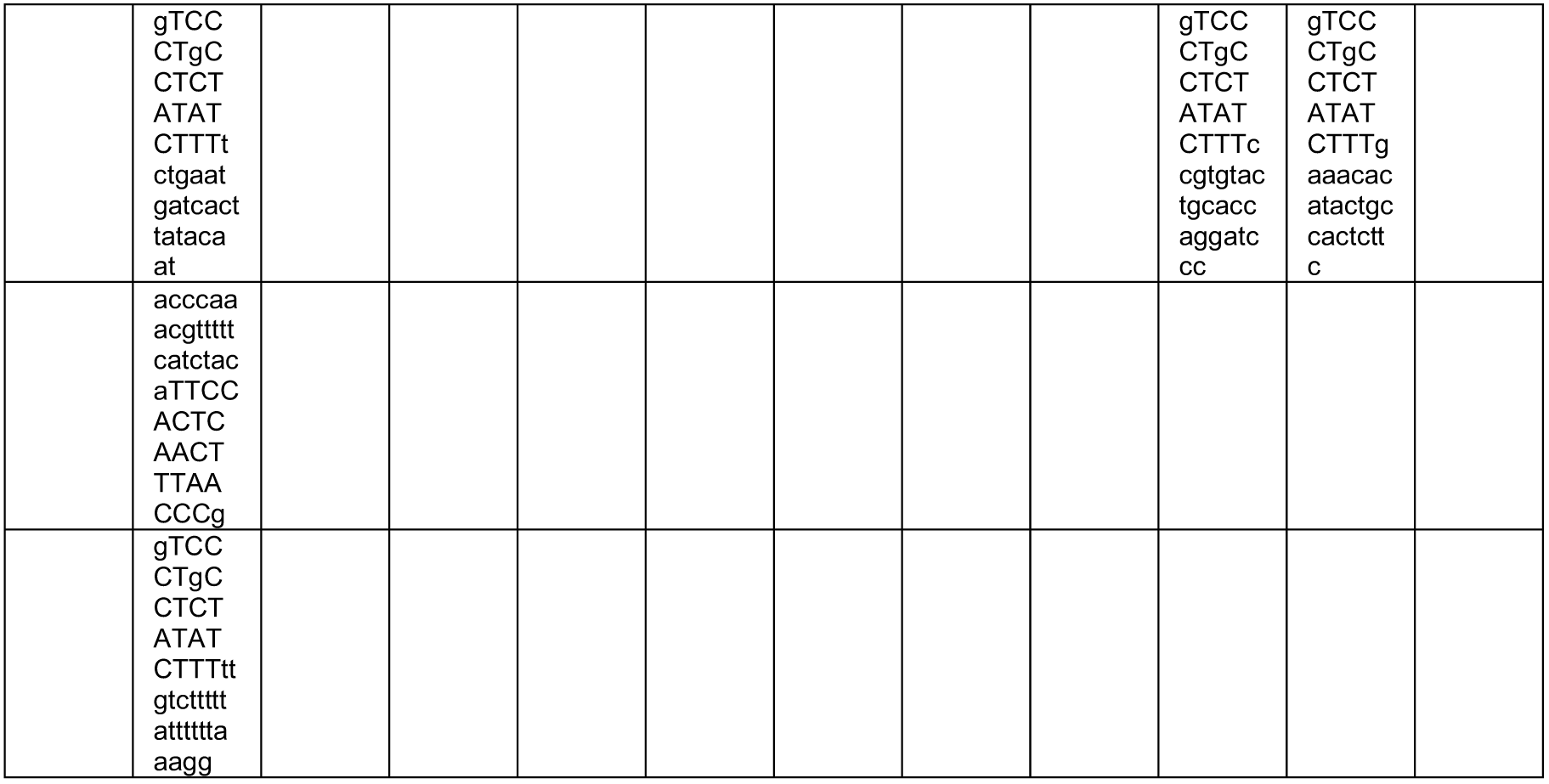
Probes for *in situ* hybridization.

## REFERENCES

Ahrens, M. B., & Engert, F. (2015). Large-scale imaging in small brains. Current opinion in neurobiology, 32, 78–86.

Ahrens, M. B., Li, J. M., Orger, M. B., Robson, D. N., Schier, A. F., Engert, F., & Portugues, R. (2012). Brain-wide neuronal dynamics during motor adaptation in zebrafish. Nature, 485(7399), 471.

Allen, W. E., Chen, M. Z., Pichamoorthy, N., Tien, R. H., Pachitariu, M., Luo, L., & Deisseroth, K. (2019). Thirst regulates motivated behavior through modulation of brainwide neural population dynamics. Science, 364(6437), 253–253.

Allen, W. E., et al. (2017). Thirst-associated preoptic neurons encode an aversive motivational drive. Science, 357(6356), 1149–1155.

Andalman, A. S., et al. (2019). Neuronal dynamics regulating brain and behavioral state transitions. Cell, 177(4), 970–985.

Aponte, Y., Atasoy, D., & Sternson, S. M. (2011). AGRP neurons are sufficient to orchestrate feeding behavior rapidly and without training. Nature neuroscience, 14(3), 351.

Betley, J. N., Cao, Z. F. H., Ritola, K. D., & Sternson, S. M. (2013). Parallel, redundant circuit organization for homeostatic control of feeding behavior. Cell, 155(6), 1337–1350.

Betley, J. N., Xu, S., Cao, Z. F. H., Gong, R., Magnus, C. J., Yu, Y., & Sternson, S. M. (2015). Neurons for hunger and thirst transmit a negative-valence teaching signal. Nature, 521(7551), 180.

Biran, J., Blechman, J., Wircer, E., & Levkowitz, G. (2018). Development and Function of the Zebrafish Neuroendocrine System. Model Animals in Neuroendocrinology: From Worm to Mouse to Man, 101–131.

Biran, J., Tahor, M., Wircer, E., & Levkowitz, G. (2015). Role of developmental factors in hypothalamic function. Frontiers in neuroanatomy, 9, 47.

Bokeh Development Team (2018). Bokeh: Python library for interactive visualization. http://www.bokeh.pydata.org.

Bourque, C. W. (2008). Central mechanisms of osmosensation and systemic osmoregulation. Nature Reviews Neuroscience, 9(7), 519.

Burdakov, D., Karnani, M. M., & Gonzalez, A. (2013). Lateral hypothalamus as a sensor-regulator in respiratory and metabolic control. Physiology & behavior, 121, 117–124.

Burnett, C. J., Li, C., Webber, E., Tsaousidou, E., Xue, S. Y., Brüning, J. C., & Krashes, M. J. (2016). Hunger-driven motivational state competition. Neuron, 92(1), 187–201.

Chiu, C. N., & Prober, D. A. (2013). Regulation of zebrafish sleep and arousal states: current and prospective approaches. Frontiers in neural circuits, 7, 58.

Choi, H. M., et al. (2018). Third-generation in situ hybridization chain reaction: multiplexed, quantitative, sensitive, versatile, robust. Development, 145(12), dev165753.

Condés-Lara, M., Rojas-Piloni, G., Martínez-Lorenzana, G., & Rodríguez-Jiménez, J. (2009). Paraventricular hypothalamic oxytocinergic cells responding to noxious stimulation and projecting to the spinal dorsal horn represent a homeostatic analgesic mechanism. European Journal of Neuroscience, 30(6), 1056–1063.

Dallman, M. F. (2005). Fast glucocorticoid actions on brain: back to the future. Frontiers in neuroendocrinology, 26(3-4), 103–108.

Dal Maschio, M., Donovan, J. C., Helmbrecht, T. O., & Baier, H. (2017). Linking neurons to network function and behavior by two-photon holographic optogenetics and volumetric imaging. Neuron, 94(4), 774–789.

Davison, J. M., et al. (2007). Transactivation from Gal4-VP16 transgenic insertions for tissue-specific cell labeling and ablation in zebrafish. Developmental biology, 304(2), 811–824.

De Marco, R. J., Groneberg, A. H., Yeh, C. M., Treviño, M., & Ryu, S. (2014). The behavior of larval zebrafish reveals stressor-mediated anorexia during early vertebrate development. Frontiers in behavioral neuroscience, 8, 367.

Dunn, T. W., Gebhardt, C., Naumann, E. A., Riegler, C., Ahrens, M. B., Engert, F., & Del Bene, F. (2016). Neural circuits underlying visually evoked escapes in larval zebrafish. Neuron, 89(3), 613–628.

Edelman, G. M., & Gally, J. A. (2001). Degeneracy and complexity in biological systems. Proceedings of the National Academy of Sciences, 98(24), 13763–13768.

Engelhard, B., et al. (2019). Specialized coding of sensory, motor and cognitive variables in VTA dopamine neurons. Nature, 570, 509–513.

Förster, D., et al. (2017). Genetic targeting and anatomical registration of neuronal populations in the zebrafish brain with a new set of BAC transgenic tools. Scientific reports, 7(1), 5230.

Freeman, J., et al. (2014). Mapping brain activity at scale with cluster computing. Nature methods, 11(9), 941.

Fujimoto, E., Stevenson, T. J., Chien, C. B., & Bonkowsky, J. L. (2011). Identification of a dopaminergic enhancer indicates complexity in vertebrate dopamine neuron phenotype specification. Developmental biology, 352(2), 393–404.

Füzesi, T., Daviu, N., Cusulin, J. I. W., Bonin, R. P., & Bains, J. S. (2016). Hypothalamic CRH neurons orchestrate complex behaviours after stress. Nature communications, 7, 11937.

Gahtan, E., Sankrithi, N., Campos, J. B., & O’Malley, D. M. (2002). Evidence for a widespread brain stem escape network in larval zebrafish. Journal of Neurophysiology, 87(1), 608–614.

Gao, Q., & Horvath, T. L. (2007). Neurobiology of feeding and energy expenditure. Annu. Rev. Neurosci., 30, 367–398.

Geerling, J. C., Shin, J. W., Chimenti, P. C., & Loewy, A. D. (2010). Paraventricular hypothalamic nucleus: axonal projections to the brainstem. Journal of Comparative Neurology, 518(9), 1460–1499.

Giovannucci, A., Friedrich, J., Gunn, P., Kalfon, J., Brown, B. L., Koay, S. A., … & Khakh, B. S. (2019). CaImAn an open source tool for scalable calcium imaging data analysis. Elife, 8, e38173.

Granger, A. J., Wallace, M. L., & Sabatini, B. L. (2017). Multi-transmitter neurons in the mammalian central nervous system. Current opinion in neurobiology, 45, 85–91.

Haesemeyer, M., Robson, D. N., Li, J. M., Schier, A. F., & Engert, F. (2015). The structure and timescales of heat perception in larval zebrafish. Cell systems, 1(5), 338–348.

Haesemeyer, M., Robson, D. N., Li, J. M., Schier, A. F., & Engert, F. (2018). A brain-wide circuit model of heat-evoked swimming behavior in larval zebrafish. Neuron, 98(4), 817–831.

Hartenstein, V. (2006). The neuroendocrine system of invertebrates: a developmental and evolutionary perspective. Journal of Endocrinology, 190(3), 555–570.

Hatta, K., Tsujii, H., & Omura, T. (2006). Cell tracking using a photoconvertible fluorescent protein. Nature protocols, 1(2), 960.

Heinrichs, S. C., & Koob, G. F. (2004). Corticotropin-releasing factor in brain: a role in activation, arousal, and affect regulation. Journal of Pharmacology and Experimental Therapeutics, 311(2), 427–440.

Herget, U., & Ryu, S. (2015). Coexpression analysis of nine neuropeptides in the neurosecretory preoptic area of larval zebrafish. Frontiers in neuroanatomy, 9, 2.

Herget, U., Wolf, A., Wullimann, M. F., & Ryu, S. (2014). Molecular neuroanatomy and chemoarchitecture of the neurosecretory preoptic-hypothalamic area in zebrafish larvae. Journal of Comparative Neurology, 522(7), 1542–1564.

Herman, J. P., & Tasker, J. G. (2016). Paraventricular hypothalamic mechanisms of chronic stress adaptation. Frontiers in endocrinology, 7, 137.

Hill, J. W. (2012). PVN pathways controlling energy homeostasis. Indian journal of endocrinology and metabolism, 16(Suppl 3), S627.

Hoshijima, K., & Hirose, S. (2007). Expression of endocrine genes in zebrafish larvae in response to environmental salinity. Journal of Endocrinology, 193(3), 481–491.

Hunter, J. D. (2007). Matplotlib: A 2D graphics environment. Computing in science & engineering, 9(3), 90.

Jennings, J. H., et al. (2015). Visualizing hypothalamic network dynamics for appetitive and consummatory behaviors. Cell, 160(3), 516–527.

Joëls, M., & Baram, T. Z. (2009). The neuro-symphony of stress. Nature reviews neuroscience, 10(6), 459.

Jones, E., Oliphant, T., & Peterson, P. (2001). SciPy: Open source scientific tools for Python.

Kato, S., et al. (2015). Global brain dynamics embed the motor command sequence of Caenorhabditis elegans. Cell, 163(3), 656–669.

Kawakami, K., et al. (2016). Gal4 driver transgenic zebrafish: powerful tools to study developmental biology, organogenesis, and neuroscience. In Advances in genetics (Vol. 95, pp. 65–87). Academic Press.

Kim, J., et al. (2019). Rapid, biphasic CRF neuronal responses encode positive and negative valence. Nature neuroscience, 22(4), 576.

Kimmel, C. B., Powell, S. L., & Metcalfe, W. K. (1982). Brain neurons which project to the spinal cord in young larvae of the zebrafish. Journal of Comparative Neurology, 205(2), 112–127.

Kimura, Y., Satou, C., & Higashijima, S. I. (2008). V2a and V2b neurons are generated by the final divisions of pair-producing progenitors in the zebrafish spinal cord. Development, 135(18), 3001–3005.

Kimura, Y., et al. (2013). Hindbrain V2a neurons in the excitation of spinal locomotor circuits during zebrafish swimming. Current Biology, 23(10), 843–849.

Kluyver, T., et al. (2016, May). Jupyter Notebooks-a publishing format for reproducible computational workflows. In ELPUB (pp. 87–90).

Korn, H., & Faber, D. S. (2005). The Mauthner cell half a century later: a neurobiological model for decision-making?. Neuron, 47(1), 13–28.

Krashes, M. J., et al. (2011). Rapid, reversible activation of AgRP neurons drives feeding behavior in mice. The Journal of clinical investigation, 121(4), 1424–1428.

Kwong, R. W., Kumai, Y., & Perry, S. F. (2014). The physiology of fish at low pH: the zebrafish as a model system. Journal of Experimental Biology, 217(5), 651–662.

Lambert, A. M., Bonkowsky, J. L., & Masino, M. A. (2012). The conserved dopaminergic diencephalospinal tract mediates vertebrate locomotor development in zebrafish larvae. Journal of Neuroscience, 32(39), 13488–13500.

Lammers, J. H. C. M., Kruk, M. R., Meelis, W., & Van der Poel, A. M. (1988). Hypothalamic substrates for brain stimulation-induced patterns of locomotion and escape jumps in the rat. Brain research, 449(1-2), 294–310.

Lee, H., et al. (2014). Scalable control of mounting and attack by Esr1+ neurons in the ventromedial hypothalamus. Nature, 509(7502), 627.

Leib, D. E., et al. (2017). The forebrain thirst circuit drives drinking through negative reinforcement. Neuron, 96(6), 1272–1281.

Li, J., et al. (2015). Intron targeting-mediated and endogenous gene integrity-maintaining knockin in zebrafish using the CRISPR/Cas9 system. Cell research, 25(5), 634.

Li, Y., et al. (2018). Hypothalamic circuits for predation and evasion. Neuron, 97(4), 911–924.

Lin, D., et al. (2011). Functional identification of an aggression locus in the mouse hypothalamus. Nature, 470(7333), 221.

Liu, N. A., et al. (2003). Pituitary corticotroph ontogeny and regulation in transgenic zebrafish. Molecular Endocrinology, 17(5), 959–966.

Löhr, H., Ryu, S., & Driever, W. (2009). Zebrafish diencephalic A11-related dopaminergic neurons share a conserved transcriptional network with neuroendocrine cell lineages. Development, 136(6), 1007–1017.

Lovett-Barron, M., Andalman, A. S., Allen, W. E., Vesuna, S., Kauvar, I., Burns, V. M., & Deisseroth, K. (2017). Ancestral circuits for the coordinated modulation of brain state. Cell, 171(6), 1411–1423.

Marder, E. (2012). Neuromodulation of neuronal circuits: back to the future. Neuron, 76(1), 1–11.

Marquart, G. D., Tabor, K. M., Bergeron, S. A., Briggman, K. L., & Burgess, H. A. (2019). Prepontine non-giant neurons drive flexible escape behavior in zebrafish. BioRxiv, 668517.

McCormick, S. D. (2001). Endocrine control of osmoregulation in teleost fish. American zoologist, 41(4), 781–794.

McCormick, S. D., & Bradshaw, D. (2006). Hormonal control of salt and water balance in vertebrates. General and comparative endocrinology, 147(1), 3–8.

McEwen, B. S. (2007). Physiology and neurobiology of stress and adaptation: central role of the brain. Physiological reviews, 87(3), 873–904.

McKinley, M. J., Denton, D. A., Ryan, P. J., Yao, S. T., Stefanidis, A., & Oldfield, B. J. (2019). From sensory circumventricular organs to cerebral cortex: neural pathways controlling thirst and hunger. Journal of neuroendocrinology, 31(3), e12689.

McKinney, W. (2010, June). Data structures for statistical computing in python. In Proceedings of the 9th Python in Science Conference (Vol. 445, pp. 51–56).

Moffitt, J. R., et al. (2018). Molecular, spatial, and functional single-cell profiling of the hypothalamic preoptic region. Science, 362(6416), eaau5324.

Mohamed, G. A., et al. (2017). Optical inhibition of larval zebrafish behaviour with anion channelrhodopsins. BMC biology, 15(1), 103.

Morrison, S. F., & Nakamura, K. (2011). Central neural pathways for thermoregulation. Frontiers in bioscience: a journal and virtual library, 16, 74.

Musall, S., Kaufman, M. T., Gluf, S., & Churchland, A. K. (2018). Movement-related activity dominates cortex during sensory-guided decision making. BioRxiv, 308288.

Nath, R. D., Chow, E. S., Wang, H., Schwarz, E. M., & Sternberg, P. W. (2016). C. elegans stress-induced sleep emerges from the collective action of multiple neuropeptides. Current Biology, 26(18), 2446–2455.

Nusbaum, M. P., Blitz, D. M., & Marder, E. (2017). Functional consequences of neuropeptide and small-molecule co-transmission. Nature Reviews Neuroscience, 18(7), 389.

Oka, Y., Butnaru, M., von Buchholtz, L., Ryba, N. J., & Zuker, C. S. (2013). High salt recruits aversive taste pathways. Nature, 494(7438), 472.

Orger, M. B., Kampff, A. R., Severi, K. E., Bollmann, J. H., & Engert, F. (2008). Control of visually guided behavior by distinct populations of spinal projection neurons. Nature neuroscience, 11(3), 327.

Parichy, D. M. (2015). The natural history of model organisms: Advancing biology through a deeper understanding of zebrafish ecology and evolution. Elife, 4, e05635.

Pedregosa, F., et al. (2011). Scikit-learn: Machine learning in Python. Journal of machine learning research, 12(Oct), 2825–2830.

Ponzio, T. A., Ni, Y., Montana, V., Parpura, V., & Hatton, G. I. (2006). Vesicular glutamate transporter expression in supraoptic neurones suggests a glutamatergic phenotype. Journal of neuroendocrinology, 18(4), 253–265.

Randlett, O., et al. (2015). Whole-brain activity mapping onto a zebrafish brain atlas. Nature methods, 12(11), 1039.

Remedios, R., Kennedy, A., Zelikowsky, M., Grewe, B. F., Schnitzer, M. J., & Anderson, D. J. (2017). Social behaviour shapes hypothalamic neural ensemble representations of conspecific sex. Nature, 550(7676), 388.

Rohlfing, T., & Maurer, C. R. (2003). Nonrigid image registration in shared-memory multiprocessor environments with application to brains, breasts, and bees. IEEE transactions on information technology in biomedicine, 7(1), 16–25.

Romanov, R. A., et al. (2017). Molecular interrogation of hypothalamic organization reveals distinct dopamine neuronal subtypes. Nature neuroscience, 20(2), 176.

Romanov, R. A., Alpár, A., Hökfelt, T., & Harkany, T. (2019). Unified Classification of Molecular, Network, and Endocrine Features of Hypothalamic Neurons. Annual review of neuroscience, 42.

Saper, C. B., & Lowell, B. B. (2014). The hypothalamus. Current Biology, 24(23), R1111–R1116.

Schöne, C., & Burdakov, D. (2012). Glutamate and GABA as rapid effectors of hypothalamic “peptidergic” neurons. Frontiers in behavioral neuroscience, 6, 81.

Schreck, C. B., & Tort, L. (2016). The concept of stress in fish. In Fish physiology (Vol. 35, pp. 1–34). Academic Press.

Schulte, P. M. (2014). What is environmental stress? Insights from fish living in a variable environment. Journal of Experimental Biology, 217(1), 23–34.

Seabold, S., & Perktold, J. (2010, June). Statsmodels: Econometric and statistical modeling with python. In Proceedings of the 9th Python in Science Conference (Vol. 57, p. 61). Scipy.

Song, K., et al. (2016). The TRPM2 channel is a hypothalamic heat sensor that limits fever and can drive hypothermia. Science, 353(6306), 1393–1398.

Spiacci Jr, A., et al. (2018). Panic-like escape response elicited in mice by exposure to CO2, but not hypoxia. Progress in Neuro-Psychopharmacology and Biological Psychiatry, 81, 178–186.

Sternson, S. M. (2013). Hypothalamic survival circuits: blueprints for purposive behaviors. Neuron, 77(5), 810–824.

Takei, Y., & Hwang, P. P. (2016). Homeostatic responses to osmotic stress. In Fish Physiology (Vol. 35, pp. 207–249). Academic Press.

Tan, C. L., & Knight, Z. A. (2018). Regulation of body temperature by the nervous system. Neuron, 98(1), 31–48.

Tan, C. L., Cooke, E. K., Leib, D. E., Lin, Y. C., Daly, G. E., Zimmerman, C. A., & Knight, Z. A. (2016). Warm-sensitive neurons that control body temperature. Cell, 167(1), 47–59.

Temizer, I., Donovan, J. C., Baier, H., & Semmelhack, J. L. (2015). A visual pathway for looming-evoked escape in larval zebrafish. Current Biology, 25(14), 1823–1834.

Tessmar-Raible, K., Raible, F., Christodoulou, F., Guy, K., Rembold, M., Hausen, H., & Arendt, D. (2007). Conserved sensory-neurosecretory cell types in annelid and fish forebrain: insights into hypothalamus evolution. Cell, 129(7), 1389–1400.

Ulrich-Lai, Y. M., & Herman, J. P. (2009). Neural regulation of endocrine and autonomic stress responses. Nature reviews neuroscience, 10(6), 397.

Van Der Walt, S., Colbert, S. C., & Varoquaux, G. (2011). The NumPy array: a structure for efficient numerical computation. Computing in Science & Engineering, 13(2), 22.

Van der Walt, S., et al. (2014). scikit-image: image processing in Python. PeerJ, 2, e453.

Vladimirov, N., et al. (2014). Light-sheet functional imaging in fictively behaving zebrafish. Nature methods, 11(9), 883.

Vladimirov, N., et al. (2018). Brain-wide circuit interrogation at the cellular level guided by online analysis of neuronal function. Nature methods, 15(12), 1117.

vom Berg-Maurer, C. M., Trivedi, C. A., Bollmann, J. H., De Marco, R. J., & Ryu, S. (2016). The severity of acute stress is represented by increased synchronous activity and recruitment of hypothalamic CRH neurons. Journal of Neuroscience, 36(11), 3350–3362.

Wang, L., Talwar, V., Osakada, T., Kuang, A., Guo, Z., Yamaguchi, T., & Lin, D. (2019). Hypothalamic control of conspecific self-defense. Cell reports, 26(7), 1747–1758.

Wang, X., et al. (2018). Three-dimensional intact-tissue sequencing of single-cell transcriptional states. Science, 361(6400), eaat5691.

Wee, C.L., et al. (2019). Zebrafish oxytocin neurons drive nocifensive behavior via brainstem premotor targets. Nature Neuroscience, doi: https://doi.org/10.1038/s41593-019-0452-x

Wen, L., et al. (2008). Visualization of monoaminergic neurons and neurotoxicity of MPTP in live transgenic zebrafish. Developmental biology, 314(1), 84–92.

Wendelaar Bonga, S. E. (1997). The stress response in fish. Physiological reviews, 77(3), 591–625.

Williams, R. H., Jensen, L. T., Verkhratsky, A., Fugger, L., & Burdakov, D. (2007). Control of hypothalamic orexin neurons by acid and CO2. Proceedings of the National Academy of Sciences, 104(25), 10685–10690.

Wingfield, J. C. (2006). Control of behavioural strategies for capricious environments. Essays in animal behaviour celebrating 50 years of Animal Behaviour, 115–133.

Wu, J. Y., Cohen, L. B., & Falk, C. X. (1994). Neuronal activity during different behaviors in Aplysia: a distributed organization?. Science, 263(5148), 820–823.

Yeh, C. M., Glöck, M., & Ryu, S. (2013). An optimized whole-body cortisol quantification method for assessing stress levels in larval zebrafish. PloS one, 8(11), e79406.

Yizhar, O., Fenno, L. E., Davidson, T. J., Mogri, M., & Deisseroth, K. (2011). Optogenetics in neural systems. Neuron, 71(1), 9–34.

Ziegler, D. R., Cullinan, W. E., & Herman, J. P. (2002). Distribution of vesicular glutamate transporter mRNA in rat hypothalamus. Journal of Comparative Neurology, 448(3), 217–229.

Zimmerman, C. A., Huey, E. L., Ahn, J. S., Beutler, L. R., Tan, C. L., Kosar, S., … & Zeng, H. (2019). A gut-to-brain signal of fluid osmolarity controls thirst satiation. Nature, 568(7750), 98.

Zimmerman, C. A., Leib, D. E., & Knight, Z. A. (2017). Neural circuits underlying thirst and fluid homeostasis. Nature Reviews Neuroscience, 18(8), 459.

